# The *Drosophila* Fab-7 boundary element modulates *Abd-B* gene activity in the genital disc by guiding an inversion of collinear chromatin organization and alternative promoter use

**DOI:** 10.1101/2022.04.26.489596

**Authors:** Laura Moniot-Perron, Benoit Moindrot, Line Manceau, Joanne Edouard, Yan Jaszczyszyn, Pascale Gilardi-Hebenstreit, Céline Hernandez, Sébastien Bloyer, Daan Noordermeer

## Abstract

*Hox* genes encode transcription factors that specify segmental identities along the Antero-Posterior body axis. These genes are organized in clusters, where their order corresponds to their activity along the body axis, an evolutionary conserved feature known as collinearity. In *Drosophila*, the BX-C cluster contains the three most posterior *Hox* genes, where their collinear activation incorporates progressive replacement of histone modifications, reorganization of 3D chromatin architecture and sequential activation of boundary elements and *cis*-regulatory regions. To dissect functional hierarchies, we compared chromatin organization in larvae and in cell lines, with a focus on the *Abd-B* gene. Our work establishes the importance of the Fab-7 boundary element for insulation between 3D domains marked by different histone modifications. Interestingly, we detected a non-canonical inversion of collinear chromatin dynamics at the *Abd-B* gene, with the active histone domain decreasing in size. This chromatin organization differentially instructed alternative *Abd-B* promoter use, thereby expanding the possibilities to regulate transcriptional output.

## Introduction

*Hox* genes encode crucial developmental regulators that specify segmental identities along the Antero-Posterior (A-P) body axis in the developing embryo of bilaterian species. A unique feature of *Hox* genes in most species is that they are organized in clusters, with their relative genomic position corresponding to their order of expression along the A-P axis (McGinnis and Krumlauf, 1992). This feature, known as collinearity, was discovered in *Drosophila melanogaster* and was later observed in other bilaterian species including mammals as well (Duboule and Dolle, 1989; Lewis, 1978). Because of this evolutionary conserved structure/function relationship, *Hox* genes have been intensively studied to decipher cluster-wide coordination of gene regulation (Duboule, 2022; Hajirnis and Mishra, 2021; Morata and Lawrence, 2022; Noordermeer and Duboule, 2013).

In *D. melanogaster*, the eight *Hox* genes are organized in two separate clusters on chromosome 3R: the Antennapedia cluster (ANT-C) and the Bithorax cluster (BX-C). The 350 kb BX-C contains the *Ubx, abd-A* and *Abd-B* genes, which specify the identity of the more posterior embryonic parasegments 5 to 14 (PS5 to PS14). To drive *Hox* gene expression in the correct parasegments (PS), the BX-C is subdivided into 10 *cis*-regulatory regions of 10 to 60 kb in size [*abx/bx, bxd/pbx* and the infra-abdominal segments 2 to 8 (*iab-2* to *iab-8*)]. Each *cis*-regulatory region contains cell type and PS-specific enhancers and is both essential and sufficient to drive the expression of its associated *Hox* gene within a single PS (Busturia and Bienz, 1993; Pirrotta, 1995; Simon *et al*., 1990).

The collinear activation of the *Hox* genes and their *cis*-regulatory elements is thought to rely on the progressive opening of the chromatin within the BX-C along the A-P axis. In this “open for business” model, the repressive H3K27me3 histone modification (associated with the Polycomb group proteins) at each *cis*-regulatory region is sequentially substituted by the active H3K4me3 modification (associated with the Trithorax group proteins) (Maeda and Karch, 2010; 2015; Schuettengruber *et al*., 2007). Consequently, in the most anterior PS5, *abx/bx* is the only *cis*-regulatory region that is marked by H3K4me3, whereas in more posterior PSs an increasingly number of *cis*-regulatory regions are covered by the H3K4me3 modification. This mechanism has been confirmed in sorted nuclei from embryonic PS4 to PS7 where the H3K27me3 mark is sequential removed from *abx/bx* up to *iab-3* (Bowman *et al*., 2014). It remains to be determined if the “open for business” model expands towards the more posterior PSs as well.

Genomic domains that carry the same histone modifications adopt a higher order 3D configuration that is known as a “contact domain” (Rao *et al*., 2014; Sexton *et al*., 2012). Within these 3D domains, intra-domain interactions are privileged over interactions with the surrounding DNA. 4C-seq and Hi-C studies in cells from *Drosophila* larvae and in cell lines have confirmed that the inactive BX-C is organized into a repressed contact domain organization that matches with the distribution of the H3K27me3 mark (Bantignies *et al*., 2011; Noordermeer and Duboule, 2013; Sexton *et al*., 2012). In mammalian cells, the sequential substitution of H3K27me3 by H3K4me3 at the *Hox* clusters along the A-P axis coincides with inactive and active contact domains of different size (Noordermeer *et al*., 2011). Similar studies in the PSs along the *Drosophila* A-P axis have not been reported, but imaging studies have identified an activity-dependent sequence of association and separation between the *Ubx* and *abd-A* genes at more anterior positions in the embryo (Cheutin and Cavalli, 2018; Mateo *et al*., 2019). In contrast, the active *Abd-B* gene does not obviously reassociate with the *Ubx* and *abd-A* genes in the posterior PSs. Likewise, comparative 3C experiments at BX-C in the *Drosophila* S2 and S3 cell lines showed that the active *Abd-B* gene and its associated *cis*-regulatory regions reduced their contacts with the inactive *Ubx* and *abd-A* genes (Lanzuolo *et al*., 2007). The unexpected behavior of the active *Abd-B* gene in these cells, where the “open for business” model predicts the absence of H3K27me3 over the nearly entire BX-C, may thus suggest that contact domains are differentially organized.

The restriction of *cis*-regulatory region activity to a single PS critically depends on the presence of Boundary Elements (BEs) between these regions (Kyrchanova *et al*., 2015). For example, the deletion of the Fab-7 boundary between *iab-6* and *iab-7* causes ectopic activation of *iab-7* in PS11, besides the normal activity of *iab-6* (Gyurkovics *et al*., 1990). Smaller deletions within Fab-7 can have an opposite effect in subsets of cells within PS11, resulting in silencing of both *iab-6* and *iab-7* (Mihaly *et al*., 1997). The likely cause for this differential outcome is the bipartite organization of Fab-7, which includes both binding sites for insulator proteins and a Polycomb and Trithorax Response Element (PRE and TRE) (Cavalli and Paro, 1998; Kyrchanova *et al*., 2015; Mihaly *et al*., 1997; Ozdemir and Gambetta, 2019). This combined function establishes the separation between neighboring *cis*-regulatory regions and provides long term maintenance of transcriptional states to those *cis*-regulatory regions as well.

How different gene regulatory layers, including *cis*-regulatory regions, histone modification domains, contact domains and BEs, functionally intersect to establish cell-type specific transcriptional programs during development remains a topic of intense interest (de Laat and Duboule, 2013; Glaser and Mundlos, 2021; Rowley and Corces, 2018; Sexton and Cavalli, 2015). Taking advantage of the detailed knowledge about the *cis*-regulatory organization of the BX-C, we have dissected functional hierarchies among gene regulatory layers at this complex, with a particular focus on the *Abd-B* gene. By combining high-resolution genomics data from larval imaginal discs and cell lines, we have confirmed that BE function extends to the insulation between histone modification domains and contact domains. Unexpectedly, in cells that represent PS12 and PS13, the organization of histone modification domains and contact domains was inversed, with the active H3K4me3 domain decreasing in size at a more posterior position. Combination of precisely calibrated and single-cell transcription analysis revealed that this inversion of collinearity differentially instructed alternative *Abd-B* promoter use, thereby providing the means for precise fine-tuning of transcriptional output.

## Results

### *Drosophila* imaginal discs and cell lines exhibit similar *Hox* gene activity patterns

The *Abd-B* gene is regulated by a 100 kb *cis*-regulatory domain that consists of 5 *iabs* that are demarcated by BEs on both sides. Transcription of *Abd-B* can be initiated from 6 alternative promoters that are located in *iab-8*, in *iab-9* or telomeric from *iab-9* (Figure 1A). To investigate how BEs, histone modification domains and 3D contact domains converge to regulate *Abd-B*, we aimed to identify cellular models where *Abd-B* is differentially expressed. Using RT-qPCR, we determined mRNA levels for *Abd-B* and five other *Hox* genes in two dissected imaginal discs from larvae and in four *in-vitro* cultured cell lines (Figure 1B). The genital disc is a mixed population of cells that originates from the most posterior PSs in the larvae, whereas the heterogeneous population of wing disc cells originate from a more anterior position (Casares *et al*., 1997).

**Figure 1:**
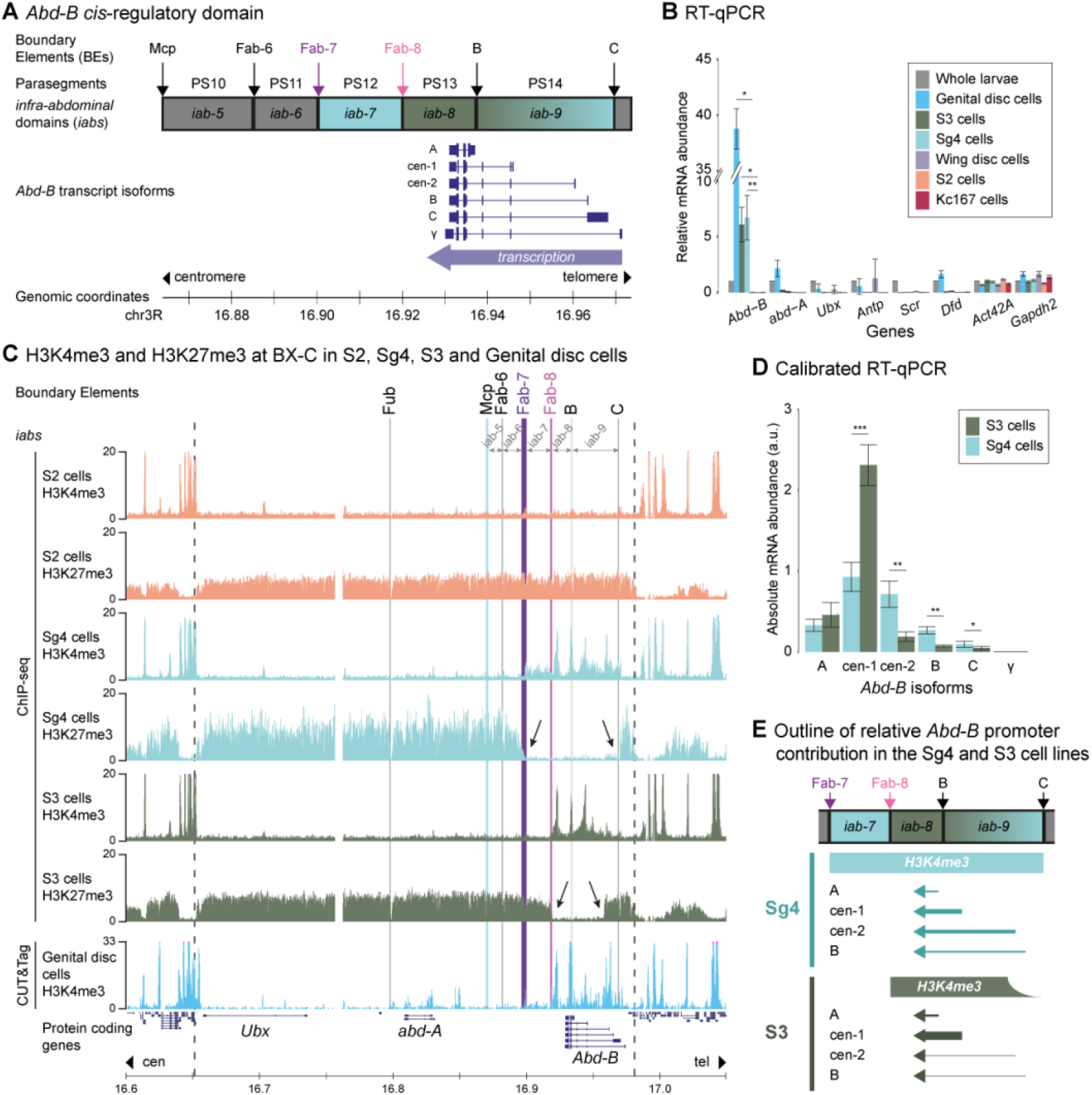
The gene expression and histone modification landscape of the BX-C in cell lines and imaginal disc cells. **A.** Overview of the ~100 kb *cis*-regulatory landscape around the *Abd-B* gene and the organization of its transcript isoforms. The *infra-abdominal* domains (*iabs*) and their associated embryonic parasegments (PS) are indicated on top, with BEs that separate *iabs* positioned in-between. The position of *Abd-B* transcript isoforms is indicated below (promoters are on the telomeric side of the gene, as indicated below). **B.** Relative *Hox* gene mRNA amounts in genital and wing disc cells and cell lines. RT-qPCR signal for each gene is normalized over the *Act42A* and *Gapdh2* housekeeping genes and expressed relative to whole larvae. Data based on biological replicates (n = 2). Error bars indicate Standard Deviation. T-test: * = p < 0.05, ** = p < 0.01. **C.** ChIP-seq data for the H3K4me3 and H3K27me3 histone marks in the S2, Sg4 and S3 cell lines and H3K4me3 CUT&Tag data in genital disc cells. The BX-C is demarcated by dashed lines, with known BEs highlighted by vertical bars. BEs and *iabs* relevant for *Abd-B* regulation are indicated on top. Location of protein coding transcripts is indicated below. **D.** Calibrated RT-qPCR to quantify the activity of alternative *Abd-B* promoters in S3 and Sg4 cells. Data are calibrated relative to a shared exon between all isoforms and to an external plasmid containing all isoform specific amplicons. Data based on biological replicates (n = 2). Error bars indicate Standard Deviation. T-test: * = p < 0.05, ** = p < 0.01, *** = p < 0.001. **E.** Schematic summary of *Abd-B* promoter contribution relative to the extent of the H3K4me3 histone modification domain in Sg4 and S3 cells. The thickness of the arrow sticks indicates the relative contribution. On top the position of BEs and *iabs* relative to the H3K4me3 histone modification domains is indicated.

Compared to whole larvae, *Abd-B* mRNA levels were robustly elevated in genital disc cells. Increased *Abd-B* amounts were detected in the Sg4 and S3 cell lines as well, albeit at lower levels as compared to the genital disc. No significant gene activity of *Abd-B* or the other *Hox* genes was detected in the wing disc or the S2 and Kc167 cell lines (Figure 1B). Our analysis thus revealed that the Sg4 and S3 cell lines mirrored the global *Hox* gene activity pattern in the genital disc, including shared *Abd-B* activity, whereas the S2 and Kc167 cell lines shared the absence of *Hox* gene expression with the wing disc.

### Histone modification domains mirror *Abd-B* expression and promoter activity, yet reveal an inversed collinear distribution

To determine how active and repressive histone modifications at the *Hox* clusters correlate with gene expression, we performed H3K4me3 and H3K27me3 ChIP-seq experiments in S2, Sg4 and S3 cells, supplemented with H3K4me3 CUT&Tag in genital discs. At the ANT-C, where no *Hox* gene activity was detected in the tested cell types, H3K4me3 was consistently depleted whereas H3K27me3 covered the entire domain (Figure S1A). In contrast, differential histone modification domains were detected at the BX-C. In S2 cells, the inactive BX-C was fully covered by H3K27me3, with no substantial peaks of H3K4me3 (Figures 1C and S1B,C). In genital discs and in Sg4 and S3 cells, the active *Abd-B* gene and its surrounding *cis*-regulatory domain were enriched for H3K4me3. Interestingly, the distribution of H3K4me3 and H3K27me3 over the active *Abd-B cis*-regulatory domain was not identical. In Sg4 cells, the entire region spanning *iab-7* to *iab-9* was devoid of H3K27me3 and enriched for H3K4me3. In contrast, in S3 cells the H3K27me3 mark was present at *iab-7* and the telomeric part of *iab-9*. Conversely, H3K4me3 was absent from *iab-7* and strongly reduced at the telomeric part of *iab-9* in these cells (Figures 1C, arrows and S1B). H3K4me3 CUT&Tag in genital disc cells suggested a mixed configuration, with robust H3K4me3 signal detected in *iab-8* and *iab-9* and a minor enrichment at *iab-7*. Activity of *Abd-B* in the different cell types was therefore accompanied by different H3K4me3 histone modification domains, which incorporated subsets of *iabs* and were demarcated by different BEs at their boundaries. Notably, in both the Sg4 and S3 cell lines the centromeric part of the BX-C, encompassing the inactive *Ubx* and *abd-A* genes and the *Abd-B-*associated *iab-5* and *iab-6*, was fully covered by H3K27me3 as well (Figure 1C). In contrast to the “open for business” model where the BX-C is thought to become sequentially devoid of the H3K27me3 mark, activity of *Abd-B* in these cells was associated with an inversed collinear presence of this repressive mark at the centromeric side of the domain.

The different domains of the H3K4me3 histone modification overlapped different sets of alternative *Abd-B* promoters, with 5 out of 6 promoters covered in Sg4 cells (A to C) and only the A and cen-1 promoters robustly marked in S3 cells (Figure 1A,C). We therefore wondered if the different cell types displayed a different preference for *Abd-B* promoter use. Calibrated RT-qPCR, with normalization to an internal *Abd-B* reference sequence, was used to quantify the relative contribution of each *Abd-B* promoter (Figure 1D). Comparison between the Sg4 and S3 cell lines revealed that the cen-1 promoter was significantly more used in S3 cells, whereas the cen-2 and B promoters contributed significantly more in Sg4 cells (Figure 1D). The combined output from these promoters is mostly similar between these two cell types though (Figure 1B). The activity of these alternative *Abd-B* promoters thus mirrored the active histone modification domains in these cell lines, yet achieving a similar transcriptional output (Figure 1E).

### The inactive BX-C is organized into two 3D contact domains

To assess the link between histone modification domains and 3D chromatin architecture at the *Hox* gene clusters, we used genome wide Hi-C and viewpoint-specific 4C-seq (Circular Chromosome Conformation Capture). We first focused on the 3D organization of the inactive clusters, which are covered by H3K27me3. By combining reanalyzed Hi-C data (Szabo *et al*.,2018) with newly generated 4C-seq data in S2 cells, we observed that the BX-C and ANT-C restricted most of their interactions within their H3K27me3-marked histone modification domains (Figures 2A and S2A,B). The inactive *Hox* clusters are therefore organized into H3K27me3-marked 3D contact domains of approximately 300-400 kb in size (Rao *et al*., 2014; Sexton *et al*., 2012). Calling of domain boundaries using the Hi-C data from S2 cells confirmed the overlap with the H3K27me3 histone modification domain, but revealed the presence of intra-domain boundaries as well: the inactive ANT-C is further divided in 3 sub-domains and the BX-C in 2 sub-domains (Figures 2A and S2A). The boundary within the BX-C is the Fub BE, which constitutes the separation between the *Ubx* and *abd-A cis-* regulatory domains (Bender and Lucas, 2013). Consequently, the inactive *abd-A* and *Abd-B cis*-regulatory domains co-occupy the same contact domain.

**Figure 2:**
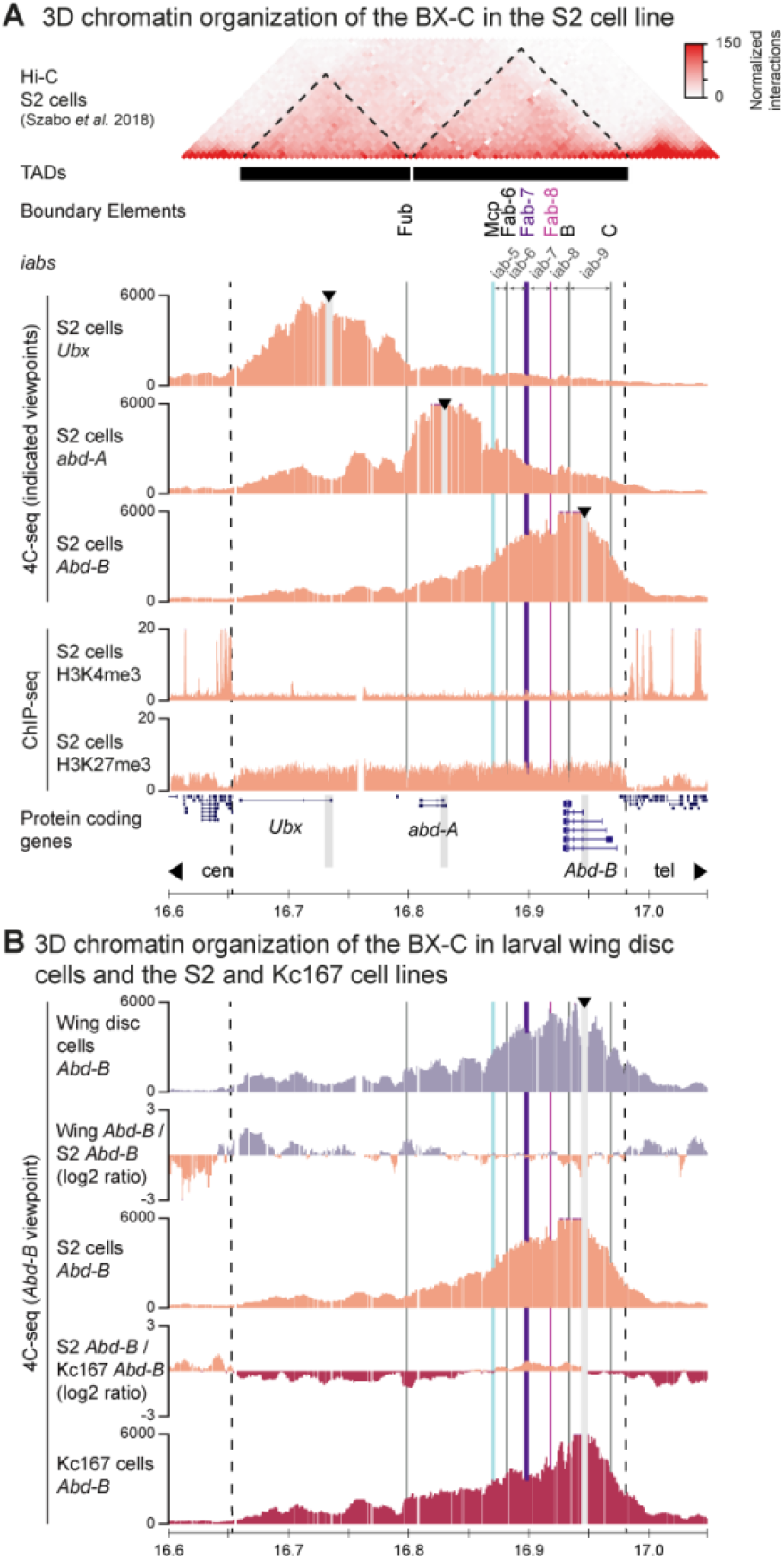
3D chromatin organization of the repressed BX-C. **A.** Hi-C (top), 4C-seq (middle) and ChIP-seq (bottom) data in the S2 cell line. Hi-C data was reanalyzed from (Szabo *et al*., 2018), with identified sub-domains indicated as black bars. 4C-seq data for viewpoints in the promoters of the *Ubx, abd-A* and *Abd-B* (cen-1) genes is indicated in-between. H3K4me3 and H3K27 ChIP-seq data is indicated below. Arrowheads indicate the positions of viewpoints. Further annotation as in Figure 1C. **B.** 4C-seq data in Wing disc cells and the S2 and Kc167 cell lines for the *Abd-B* cen-1 promoter viewpoint. In-between, the log2 ratio of interactions is shown.

Next, we used 4C-seq to address if the same contact domains were present at the inactive ANT-C and BX-C in larval Wing discs and the Kc167 cell line as well. 4C-seq interaction patterns in these cell types were highly similar to those in S2 cells, confirming that the inactive *Hox* clusters adopt a similar 3D contact domain organization (Figures 2B and S2C,D).

### Active *Abd-B* dissociates from the repressed BX-C contact domains in Sg4 cells

To determine the 3D chromatin organization of the *Hox* clusters when *Abd-B* is active, we first performed Hi-C and 4C-seq in the Sg4 cell line. The contact domain organization of the inactive ANT-C in Sg4 cells was comparable to the previously established organization in S2 cells, confirming that interactions were not drastically reorganized between these cell types (Figures S2A and S3A,B). In contrast, the 3D organization of the BX-C was markedly different between Sg4 and S2 cells. Our Hi-C data revealed that the BX-C in Sg4 cells is organized into 3 sub-domains, with each *Hox* gene occupying its own domain (Figure 3A). Whereas the boundary between the repressed *Ubx* and *abd-A* genes remained at the Fub BE, a new boundary had appeared that coincided with the Fab-7 BE element. 4C-seq experiments confirmed this observation, which was particularly visible upon comparison of *Abd-B* promoter interactions between the Sg4 and S2 cell lines. Indeed, the Fab-7 BE acted as the boundary between the *Abd-B* H3K4me3 histone modification domain and the repressed contact domain consisting of *iab-5, iab-6* and the *abd-A cis*-regulatory domains, with a strong loss of interactions in Sg4 cells observed in the latter domain (Figure 3A; yellow shading). We thus concluded that the *Abd-B* H3K4me3 histone modification domain, bordered by the Fab-7 and C BEs and comprising *iab-7* to *iab-9*, was organized as an active contact domain that was physically dissociated from the inactive *abd-A* contact domain.

**Figure 3:**
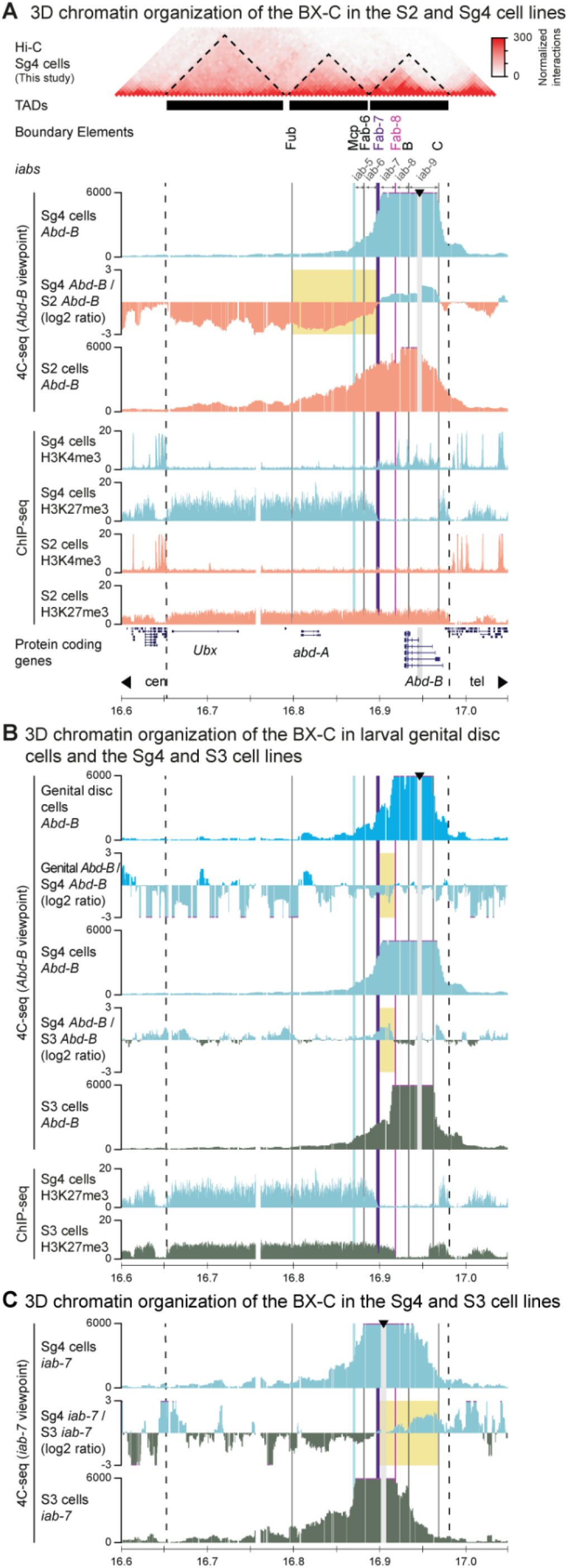
3D chromatin organization of the BX-C in cell types where *Abd-B* is active. **A.** Hi-C (top) and 4C-seq (middle) and ChIP-seq (bottom) data in the Sg4 cell line. Sub-domains as identified from Hi-C are indicated as black bars. 4C-seq data for the *Abd-B* cen-1 promoter viewpoint in S2 (*Abd-B* inactive) and Sg4 cell lines and the log2 ratio of interactions is indicated in-between. The yellow rectangle highlights the domain of reduced interactions in Sg4 cells whose border coincides with the Fab-7 BE. H3K4me3 and H3K27 ChIP-seq data in the S2 and Sg4 cell lines are indicated below. Arrowheads indicate the position of the viewpoint. Further annotation as in Figure 1C. **B.** 4C-seq data in genital disc cells and the Sg4 and S3 lines for the *Abd-B* cen-1 promoter viewpoint. In-between, the log2 ratio of interactions is shown. Yellow rectangles highlight the domains of increased interactions in Sg4 cells that cover *iab-7*. **C.** 4C-seq data in the Sg4 and S3 cell lines for *iab-7* viewpoint. In-between, the log2 ratio of interactions is shown. The yellow rectangle highlights the domain of increased interactions in Sg4 cells that covers *iab-9*.

### The active *Abd-B* 3D contact domain is bordered by different BEs in different cell types

Next, we wondered if the same contact domain organization was present in cells from the larval genital disc and the S3 cell line, where the *Abd-B* gene is active but associated with H3K4me3 histone modification domains of different size (Figure 1B,C). 4C-seq at the inactive ANT-C revealed a mostly comparable 3D chromatin organization in all cell types, confirming that global 3D interactions were similar between cell types (Figure S3C). Comparison of interactions further confirmed the dissociation of *Abd-B* from the inactive remainder of the BX-C, but also revealed noticeable differences in 3D contact domain boundaries (Figure 3B). Whereas in Sg4 cells a steep drop in interactions coincided with the Fab-7 BE, a similar drop in S3 cells coincided with the Fab-8 BE. As a result, interactions of the *Abd-B* cen-1 promoter viewpoint with *iab-7* where considerably depleted in the S3 cell line as compared to Sg4 cells. Conversely, the inactive *Ubx* and *abd-A* promoters contacted *iab-7* more in the S3 cells (Figure 3B; yellow shading and Figure S3D). This observation was further confirmed using a viewpoint at *iab-7* itself, which associated more with the active *iab-8* and *iab-9* in Sg4 cells and with the centromeric inactive part of the BX-C in S3 cells (Figures 3C; yellow shading). The use of different BEs in the different cell lines thus mirrored the boundaries of the different H3K4me3 histone modification domains (Figure 1C) and the association of *iab-7* with the active or inactive contact domain directly coincided with the presence of the H3K4me3 or H3K27me3 histone modifications.

A similar difference was observed for the *Abd-B* cen-1 promoter viewpoint when comparing the genital disc cells to the Sg4 cell line, although the heterogeneous nature of the genital disc provided less discrete patterns of interactions. In genital disc cells, a steep drop in interactions coincided with the Fab-8 BE (like in S3 cells), yet a minor drop at the Fab-7 BE was observed as well (like in Sg4 cells). Both BEs thus appear to contribute to the separation between the active and inactive contact domain, with *iab-7* differentially associating with either domain (Figure 3B; yellow shading).

Based on the similarities between BE use, histone modification and contact domains and knowledge about PS-specific activity of BEs and *iabs* (Kyrchanova *et al*., 2015), we considered it likely that the differences in BX-C organization between the Sg4 and S3 cell lines mirrored different chromatin configurations that were present within the genital discs. Specifically, Sg4 cells resembled cells that represented PS12, where the Fab-7 BE acted as boundary between the centromeric inactive contact domain and the active H3K4me3 contact domain that spanned *iab-7* to *iab-9*. In contrast, S3 cells resembled cells from PS13, where the Fab-8 BE constituted the boundary between the centromeric inactive contact domain and the active H3K4me3 contact domain that spanned the *iab-8* and to a lesser extent *iab-9*. Our data in these cell lines thus provided valuable snapshots of chromatin organization that mirrored the organization in different PSs. These observations confirmed the direct overlap between histone modification domains and 3D contact domains and identified BEs as the DNA elements that provide cell-type specific separation between the 3D histone modification domains of varying size.

### Absence of the Fab-7 BE reduces overall *Abd-B* activity in genital discs

To directly assess the importance of BE function for separation of 3D histone modification domains in a relevant developmental context, we analyzed an existing *Drosophila* mutant line where the Fab-7 element is absent (Gyurkovics *et al*., 1990). In this Fab-7^1^ strain, the absence of a 4.3 kb DNA fragment that encompasses both its BE and PRE function causes the ectopic activation of *iab-7* in the more anterior abdominal segments of the adult fly (Gyurkovics *et al*., 1990; Karch *et al*., 1994; Singh and Mishra, 2015). To determine how the Fab-7 BE impacts on global *Hox* gene expression in larvae, we performed RT-qPCR in genital and wing disc cells from the WT and homozygous Fab-7^1^ strains. The larval genital disc is thought to encompass the most posterior PSs (from PS13), whereas our results suggested a contribution from PS12 as well [Figure 3B and (Kyrchanova *et al*., 2015; Sanchez and Guerrero, 2001)]. Upon removal of Fab-7 all *Hox* genes remained inactive in wing disc cells, yet in genital disc cells we observed a 10% reduction in *Abd-B* mRNA abundance (p < 0.06) (Figure 4A). The Fab-7 BE, which borders *iab-7* that is active in PS12, therefore moderately contributes to overall *Abd-B* activity in the genital disc.

**Figure 4:**
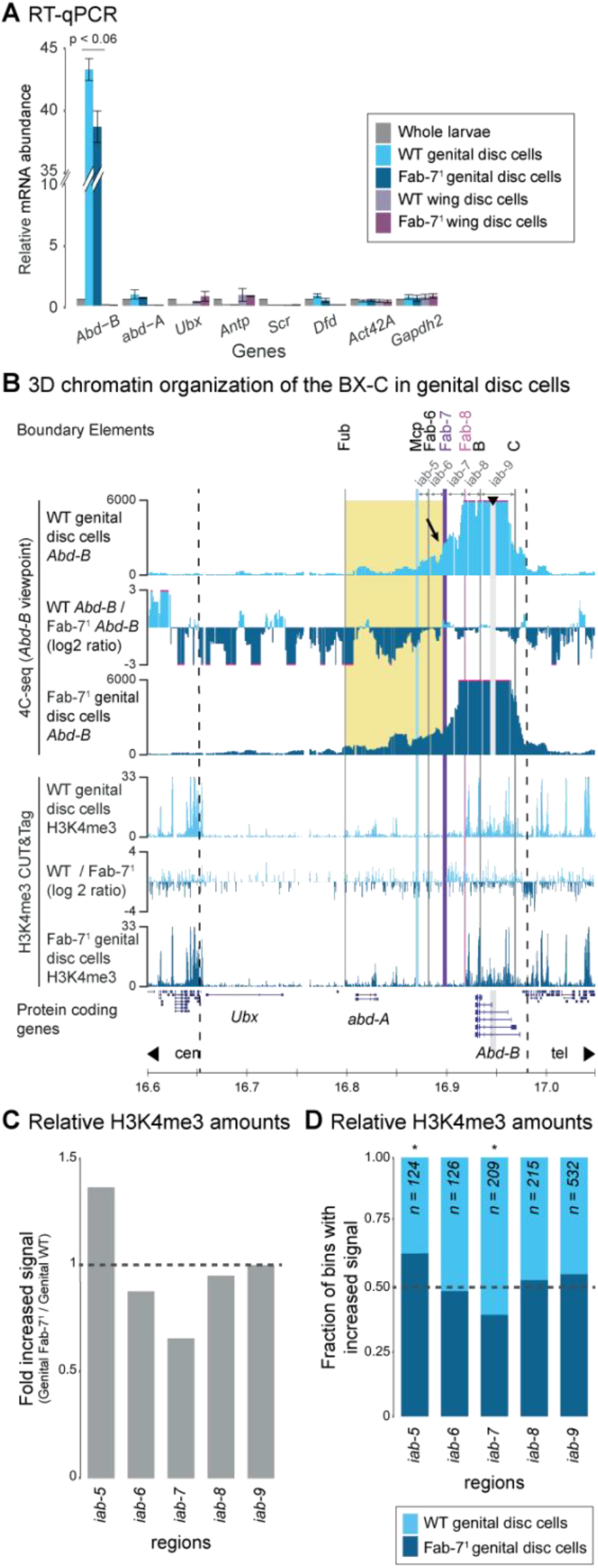
*Hox* gene expression and 3D chromatin organization in WT and Fab-7^1^ deletion larval imaginal disc cells. **A.** Relative *Hox* gene mRNA amounts in WT and Fab-7^1^ deletion wing and genital disc cells (blue). RT-qPCR signal for each gene is normalized over the *Act42A* and *Gapdh2* housekeeping genes and normalized relative to WT whole larvae. Data based on biological replicates (n = 2). Error bars indicate Standard Deviation. **B.** 4C-seq (top) and H3K4me3 CUT&Tag (bottom) data in WT and Fab-7^1^ deletion genital disc cells. 4C-seq data is shown for the *Abd-B* cen-1 promoter viewpoint, with the log2 ratio of interactions indicated in-between. The abrupt drop of interactions at the Fab-7 BE in WT cells is highlighted with the arrow and the region with increased interactions over the *abd-A* gene in Fab-7^1^ deletion cells is highlighted with the yellow rectangle. The log2 ratio of CUT&Tag signal (50 bp bins) is indicated in-between the tracks. Further annotation as in Figure 1C. **C.** Ratio of total H3K4me3 CUT&Tag signal in the indicated domains. Values > 1 indicate a gain of domain-wide signal in Fab-7^1^ deletion cells and values < 1 indicate a loss of domain-wide signal. **D.** Fraction of H3K4me3 CUT&Tag bins (50 bp windows, normalized signal) with increased signal in WT or Fab-7^1^ deletion cells. G-test: *: p < 0.05.

### Absence of Fab-7 reorganizes contact domains and histone modification domains

Next, we used 4C-seq to determine if the absence of the Fab-7 BE had a noticeable effect on 3D contact domain structure. In wing disc cells, where *Abd-B* is repressed, we detected no notable differences between the WT and deletion strain (Figure S4A). In genital disc cells considerable differences were observed though. Whereas in the absence of Fab-7 the steep drop in interactions at the Fab-8 BE remained (pink line), the smaller drop that was observed at the WT Fab-7 BE (purple line) had disappeared (Figure 4B). Moreover, in the absence of Fab-7 BE, the interactions of the *Abd-B* viewpoint were globally increased over the entire BX-C, with a particularly prominent signal now detected within the *abd-A cis*-regulatory domain and beyond (Figure 4B; yellow shading). Conversely, the inactive *Ubx* and *abd-A* viewpoints increased their contacts with the *Abd-B* regulatory domain, which was particularly prominent at *iab-7* (Figure S4B). Based on the observed patterns in the Sg4 and S3 cell lines, we envision that the separation between the active and inactive contact domains at the Fab-7 BE may have been lost in the subset of PS12 cells within the mixed genital disc cell population (Figure 3B). The loss of separation between the active and inactive contact domains creates an organization that resembles the S2 cell line (Figures 2A and 3A). This increased association with the repressive 3D contact domain in a subset of genital disc cells may explain the observed moderate reduction in *Abd-B* mRNA levels (Figure 4A).

To determine if the increased association with the inactive contact domain had an influence on the H3K4me3 histone modification domain, we generated H3K4me3 CUT&Tag in Fab-7 deletion genital discs (Figure 4B, bottom tracks). Comparison of H3K4me3 distribution to WT cells revealed a significant 1.5-fold reduction of this active mark within *iab-7*. In contrast, the mark was significantly increased within *iab-5*, although the overall signal remained low within this domain. Within the mixed population of the genital disc cells, no significant difference was detected at *iab-8* and *iab-9*, where the majority of *Abd-B* promoters are localized (Figure 4C,D). The reduced H3K4me3 levels at the *iab-7* regulatory domain therefore emerged as the most apparent link with the globally decreased *Abd-B* expression. As both Fab-7 and *iab-7* activity are associated with PS12, we further envisioned that the increased association with the repressive 3D contact domain in the absence of this BE may be causally linked to the loss of H3K4me3 in this cell population.

### Absence of Fab-7 reorganizes *Abd-B* promoter choice in genital disc cells

Next, we wondered if the observed reduction of H3K4me3 at *iab-7* in the absence of the Fab-7 BE could be associated with a differential choice of alternative *Abd-B* promoters, similar to the differences as observed in the Sg4 and S3 cell lines (Figure 1D). To address this question, we first assessed *Abd-B* promoter choice using calibrated RT-qPCR in WT and Fab-7^1^ genital disc cells (Figure 5A). Confirming our previous observation that *Abd-B* activity was reduced in the absence of Fab-7, we noticed that mRNA levels for most *Abd-B* isoforms were less abundant as well (Figures 4A and 5A). Although some promoter specificity could be observed, the pan-cellular nature of this analysis precluded the distinction between a global reduction or reduced numbers of *Abd-B* expressing cells.

**Figure 5:**
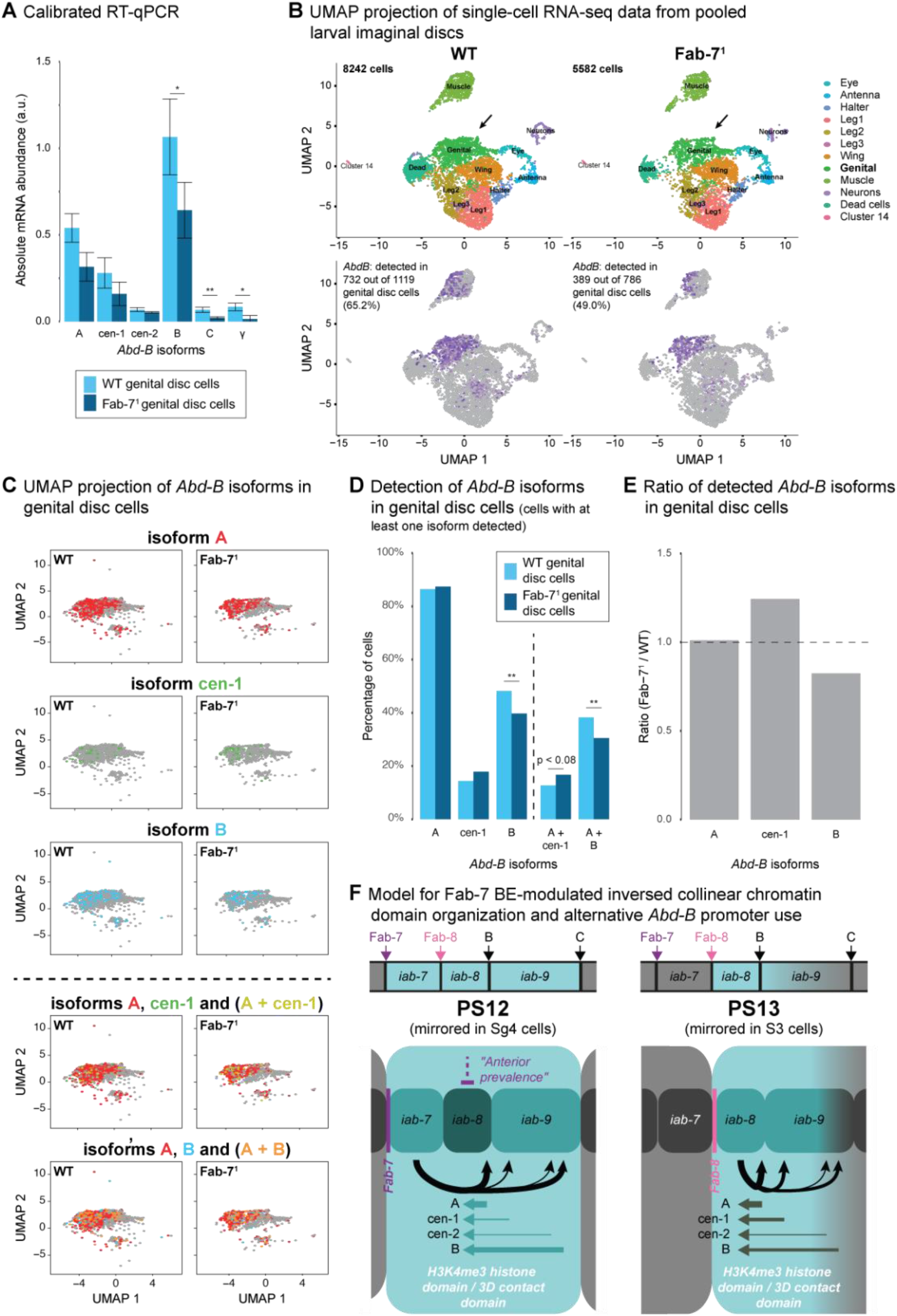
*Abd-B* promoter activity in WT and Fab-7^1^ deletion imaginal disc cells. **A.** Calibrated RT-qPCR to quantify the activity of alternative *Abd-B* promoters in WT and Fab-7 deletion genital disc cells. Data are calibrated relative to a shared exons between all isoforms and to an external plasmid containing all isoform specific amplificons. Data based on biological replicates (n = 2). Error bars indicate Standard Deviation. **B.** UMAP projections of integrated single-cell RNA-seq data from pools of WT and Fab-7 deletion imaginal disc cells. On the top, the identified clusters are indicated, which comprises clusters for the 8 different discs and 4 other clusters (with data separated for WT (left) and Fab-7 deletion (right) cells). Arrow highlights the merged genital disc cluster. On the bottom, the presence of *Abd-B* mRNA (purple) within the pools of imaginal disc cells is indicated. **C.** UMAP projections of genital disc cells for the indicated *Abd-B* promoters (top panels) and pairs of promoters (bottom panels). Color-coding as indicated in the header of each panel. **D.** Histogram showing the percentage of cells where mRNA from the indicated promoters or pairs of promoters is detected (percentages within the population of genital disc cells where at least one isoform of *Abd*-*B* is detected). G-test: ** p < 0.01. **E.** Histogram showing the ratio of detected *Abd*-*B* mRNA (Fab-7^1^ deletion / WT genital disc cells) originating from the A, cen-1 and B promoters. **F.** Model for chromatin organization and *Abd-B* promoter use is PS12 and PS13. The light blue spheres indicate the 3D contact domains, with *iabs* schematically positioned within. Colored arrows indicate alternative *Abd-B* transcripts and black arrows indicate transcriptionally activating influence, with thickness of the arrow sticks indicating relative contribution. On top the position of BEs and *iabs* relative to the H3K4me3 histone modification domains is indicated.

To overcome this limitation, we performed single-cell RNA-seq analysis on pools of eight dissected larval imaginal discs using a setup that allowed the identification of promoter origin. After filtering of low-quality cells, we obtained data from 8242 WT cells and 5582 Fab-7 deletion cells (Figure 5B, top). Although fewer Fab-7^1^ cells were recovered, similar numbers of unique mRNA molecules and genes were detected in both cell types, thus confirming similar data quality (Figure S5A,B). Clustering of the single-cell RNA-seq data, obtained from the combined WT and Fab-7^1^ deletion pools of imaginal discs, identified 14 distinct clusters of cells. Based on tissues-specific marker genes, including the *Hox* genes, we traced back the eight different larval imaginal discs and four other cell population that were inadvertently included in our dissection procedure (Figures 5B and S5C). Both genital disc and muscle cells could be traced back to two clusters, which we merged in the downstream analysis. Importantly, the 12 remaining clusters all contained a mix of WT and Fab-7^1^ deletion cells, confirming that the absence of the Fab-7 BE did not influence cell and cluster identity (Figure 5B).

*Abd-B* transcripts, originating from any promoter, were primarily detected in the genital disc and in a sub-population of muscle cell clusters, both in the WT and Fab-7^1^ deletion cells (Figure 5B). We restricted the remainder of our analysis on the cells in the genital disc cluster, which was comprised of 1119 WT and 786 Fab-7^1^ cells. We could detect *Abd-B* transcripts in 65% of WT cells, yet this was reduced to 49% of Fab-7 deletion cells (Figures 5B and S5D; despite similar single-cell RNA-seq quality metrics). The reduction of *Abd-B* expression levels in Fab-7^1^ cells, as detected by RT-qPCR, may thus (partially) be explained by the lower number of cells where *Abd-B* was robustly expressed as detected in our single-cell RNAs-seq analysis (Figures 4A and 5B).

Notably, we could detect mRNA from all other *Hox* genes in smaller numbers of genital disc cells as well, showing that the repression of *Hox* genes in the larval genital disc is not complete (Figure S5C,D). Similar observations were made in wing disc cells and in a limited single-cell RNA-seq analysis of the Sg4 and S2 cell lines, where *abd-A* mRNA was detected in a subset of Sg4 cells as well (Figure S5D,E).

Next, we quantitatively assessed the presence of *Abd-B* transcripts originating from different promoters. For this purpose, we focused only on cells where at least one *Abd-B* isoform was detected. mRNA originating from the A, cen-1 and B promoters was present in the largest numbers of individual cells, which was in line with our calibrated RT-qPCR result (Figures 5A and S5F). By focusing on these three promoters, we identified differences in the response to Fab-7 removal (Figure 5C-E). Whereas the relative number of cells containing mRNA from the A promoter remained similar, the number of cells that were positive for the cen-1 and B promoters showed an opposing dynamic. Cells carrying mRNA from the cen-1 promoter were relatively more abundant upon the deletion of the Fab-7 BE, whereas relatively more WT cells carried mRNA from the B promoter (Figure 5D-E). This observation was further strengthened by limiting the analysis to cells where mRNA from multiple promoters was detected. Indeed, the combination of mRNA from the A and more centromeric cen-1 promoters was enriched upon Fab-7 deletion and the combination of the A and more telomeric B promoters was enriched in WT cells (Figure 5D-E). Our single-cell RNA-seq data thus revealed that in the absence of the Fab-7 BE, both the number of *Abd-B* expressing cells was reduced and that the *Abd-B* promoter use was reorganized. mRNA originating from the cen-1 promoter, located at a centromeric position within *iab-9* and privileged by the chromatin organization in PS13, became relatively more abundant. Conversely, mRNA from the more telomeric B promoter, associated with the chromatin organization in PS12, had a reduced presence. The analysis of single-cell promoter use thus reinforced the notion that fewer cells from PS12 contributed to the detected *Abd-B* activity in the genital discs.

## Discussion

In this study we determined how histone modification domains, 3D chromatin organization and BEs functionally engage at the *Abd-B* regulatory domain in cells that represent different parasegments. In cells where the BX-C was fully repressed, most of its chromatin interactions precisely overlapped with the presence of the H3K27me3 histone mark, confirming previous observations that the H3K27me3 histone modification domain and the 3D contact domain are tightly linked (Lanzuolo *et al*., 2007; Mateo *et al*., 2019). In cells where *Abd-B* was active, the H3K4me3-marked *cis*-regulatory domain was organized into a contact domain that dissociated from the inactive domain. BEs on both sides of the active contact domain acted as boundaries, providing a new mechanistic detail to previous observations that active and inactive genes and domains within the BX-C are physically separated (Cheutin and Cavalli, 2018; Lanzuolo *et al*., 2007; Mateo *et al*., 2019). Different BEs could act as boundary though, with either Fab-7 or Fab-8 separating the histone modification and contact domains. We argue that these different chromatin organizations represent the organization in PS12 and PS13, which both contribute to the heterogeneous genital disc cell population (Figure 5F). Like at mammalian *Hox* gene clusters, active and inactive contact domains of different size are therefore present along the *Drosophila* A-P axis (Noordermeer and Duboule, 2013; Noordermeer *et al*., 2011). Absence of the Fab-7 BE in the heterogeneous genital disc cell population coincided with a loss of H3K4me3 at *iab-7* in a subpopulation of cells, which we presume were cells that originated from PS12 where Fab-7 functions as active BE. This result establishes the instructive nature of BEs in demarcating histone modification and contact domains. The loss of H3K4me3 over *iab-7* appears contrary to the outcome in larval PS11 though, where the absence of Fab-7 causes ectopic activation of *iab-7* (Gyurkovics *et al*., 1990). Upon smaller deletions of Fab-7 in PS11, a crosstalk between PREs and TREs could induce silencing of *iab-6* and *iab-7* in subsets of cell though (Mihaly *et al*., 1997). PS12-specific crosstalk between PREs and TREs in *Fab-6* and *Fab-8* may therefore explain the silencing of *iab-7* in these cells as well (Kyrchanova *et al*., 2015).

The chromatin organization of the active domain in PS12 and PS13 covered different numbers of alternative *Abd-B* promoters. As a result, BE use influenced the abundance of *Abd-B* isoforms (Figure 5F). In line with the notion that a single *iab* is responsible for the activation of *Abd-B* in a single PS, we did not detect a large difference in the combined amounts of *Abd-B* transcripts in the Sg4 and S3 cell lines. This raises the question why alternative *Abd-B* promoters are differentially modulated by *iab-7* and *iab-8*. Although not fully consistent with the currently established promoters, previous studies have suggested that the different mRNA molecules that originate from the cen-1, cen-2, B and C promoters encode the same Abd-B protein isoform [Figure 1A and (Celniker *et al*., 1989; Zavortink and Sakonju, 1989)]. Although we can’t rule out that the alternative transcripts have a direct effect on protein abundance (e.g., through different mRNA stability or translation initiation kinetics), another explanation may be that the different promoters provide a means for fine-tuning transcriptional output in the different PSs, either by increasing the total number of transcription initiation events or by buffering against inherently different promoter affinities between enhancers located in *iab-7* and *iab-8*.

According to the “open for business” model for BX-C activation, the H3K27me3 mark should be progressively replaced by the active H3K4me3 mark along the A-P axis (Maeda and Karch, 2010; 2015). Our description of the chromatin organization associated with the previously uncharacterized PS12 and PS13 has identified an unexpected inversion of collinear histone modifications at the *Abd-B* regulatory domain. Not only was the centromeric part of the BX-C, including the *Abd-B* regulating *iab-5* and *iab-6*, covered by the H3K27me3 mark in PS12, but in S3 cells— representative of the more posterior PS13—this repressive domain had increased in size to included *iab-7* as well (Figure 5F). Although we can’t formally rule out that this inversion of collinearity is unique to larval cells, we consider this unlikely. First, the cell lines included in our study are of embryonic origin and second, our result provides an improved context for previous observations that *Abd-B* does not reassociate with the *Ubx* and *abd-A* genes in the most posterior cell types of the *Drosophila* embryo (Cheutin and Cavalli, 2018; Mateo *et al*., 2019). Intriguingly, in the absence of Fab-7, we and others detected increased interactions of *Abd-B* with the supposedly H3K27me3-marked centromeric part of BX-C [Figure 4B and (Mateo *et al*., 2019)]. Combined with the considerable reduction in the number of genital disc cells where *Abd-B* mRNA is detected and the supposed restriction of Fab-7 activity in PS12, we hypothesize that the BX-C had adopted an inactive chromatin configuration in this cell population, which was reminiscent of the 3D organization in wing disc cells and the S2 and Kc167 cell lines.

The non-canonical inversion of collinearity in the most posterior PSs may relate to the genetic organization of the *Abd-B cis*-regulatory domain. Whereas the promoters of the *Ubx* and *abd-A* genes are located in the most centromeric *cis*-regulatory regions for each gene, the promoters of *Abd-B* are located in the most telomeric *iab-8* and *iab-9* (Figure 1A). Instead of a sequential activation of *cis*-regulatory regions towards the telomeric side, as observed for *Ubx* and *abd-A*, the activation of *Abd-B* by the more centromeric *iabs* may critically depend on the absence of H3K27me3 from *iab-8* and *iab-9* as well. PS-specific regulation by a single *iab* is subsequently achieved by two repressive mechanisms that maintain the inactive state of enhancers in the other *iabs*. First, the more anterior *iabs* remain inactive through the inversion of collinear H3K27me3-mediated repression, as is the case for *iab-5* and *iab-6* in PS12 and *iab-5* to *iab7* in PS13. Second, the enhancers in the more telomeric *iabs*, which are included in the H3K4me3-marked contact domain, are kept in an inactive state through an “anterior prevalence” of the most centromeric *iab* within the H3K4me3-marked contact domain. This is particularly relevant for the enhancers in *iab-8* in both PS12 cells and the Sg4 cell line, although the mechanistic underpinning of this “anterior prevalence” remains to be confirmed (Figure 5F). Similarly, it remains to be determined if *Abd-B* activation by *iab-5* and *iab-6* in the more anterior PS10 and PS11, which are not part of the genitalia, involves an inversion of collinear chromatin organization and “anterior prevalence” over *iab-6* to *iab-8* as well.

Interestingly, a non-canonical inversion of collinear chromatin dynamics is observed at the mammalian *HoxA* and *HoxD* clusters during the development of the mammalian genitalia and digits. At these clusters, the *Hox* genes that are expressed at the most anterior positions along the A-P axis are, within the genitalia and digits, covered by the repressive H3K27me3 modification (Lonfat et al., 2014; Montavon et al., 2011). Instead, genes from the mammalian “posterior” group-9 to group-13 *Hox* genes are active, which are orthologous genes of *Abd-B* that arose through serial duplications of a shared ancestor (Izpisua-Belmonte et al., 1991). Like in *Drosophila*, the activation of mammalian *Hox* genes in the genitalia and digits requires dynamic contacts with distant enhancers (Lonfat *et al*., 2014; Montavon *et al*., 2011). Although the organization of the *cis*-regulatory landscapes have considerably diverged between these evolutionary distant species, this raises the intriguing possibility that the inversion of collinear chromatin organization involving *Abd-B* class genes is an ancestral feature, possibly associated with the emergence of genitalia. The investigation of the collinear activation of *Abd-B* group genes in the genitalia of other distant bilaterian lineages may confirm if the inversion of collinearity is indeed an evolutionary conserved feature that has been inherited from a common ancestor.

## Acknowledgments

We thank Sylvina Bouleau, Emilie Brasset, Fabienne Malagnac and the members of the Noordermeer lab for useful discussion and we thank Denis Duboule and Maria Cristina Gambetta for critical reading of the manuscript. We are grateful to François Karch, Jacques Montagne, Vanessa Ribes and Matthieu Sanial for sharing of reagents and materials. We thank the I2BC High throughput Sequencing Platform for sequencing experiments and single-cell sequencing library preparation. Work in the Noordermeer lab is supported by funds from the Agence Nationale de la Recherche (ANR-14-ACHN-0009-01, ANR-16-TERC-0027-01, ANR-17-CE12-0001-02, ANR-18-CE12-0022-02 and ANR-21-CE12-0034-01), PlanCancer (19CS145-00) and the Fondation Bettencourt Schueller to D.N. L.M.-P. was supported by thesis grants from the Fondation pour la Recherche Médicale (FDT202012010470) and the Université Paris-Saclay (Ecole Doctorale Structure et Dynamique des Systèmes Vivants). L.M. received thesis funding from the Institut Jacques Monod. The funders had no influence on the design of the project or the decision to publish.

## Author contributions

Conceptualization, S.B. and D.N.; Methodology, L.M.-P., B.M., S.B. and D.N.; Formal analysis, L.M.-P., B.M., C.H. and D.N.; Investigation, L.M.-P., B.M., L. M., J.E., Y.J. and P.G.-H.; Writing – Original Draft, L.M.-P., S.B. and D.N.; Writing – Review & Editing, L.M.-P., B.M., S.B. and D.N.; Supervision, S.B. and D.N.

## Declaration of interests

The authors declare no competing interests.

## Data availability

All unprocessed Illumina sequencing data generated in this study have been deposited in the European Nucleotide Archive (EMBL-EBI ENA) under accession number PRJEB52393.

## Materials and methods

### Fly stocks

WT flies (w^1118^ background) were a gift from Jacques Montagne (I2BC, France) and Fab7^1^ deletion flies were a gift from François Karch (University of Geneva, Switzerland). In Fab7^1^ flies, a 4.3 kb region between *Abd-B* and *abd-A* that contains the Fab-7 element is absent (Gyurkovics *et al*., 1990). Fly stocks were maintained and cultured using standard cornmeal yeast extract medium at 25°C.

### Cell culture

S2, Sg4 and S3 cells were ordered from Drosophila Genomics Resource Center (Indiana University, USA). Kc167 cells were a kind gift from Matthieu Sanial (IJM, France). S2 cells were grown in Schneider medium (ThermoFisher Scientific), Sg4, S3 and Kc167 cells were grown in M3+BPYE medium (Sigma-Aldrich). Culture mediums were supplemented with 1% penicillin/streptomycin (ThermoFisher Scientific) and different amounts of Fetal Bovine Serum (FBS): 10% for S2 cells; 12.5% for the Sg4, S3 and Kc167 cells). Conventional Gibco FBS was used (ThermoFisher Scientific) for S2 and S3 cells, whereas Gibco Performance FBS was used for Sg4 and Kc167 cells. Cells were inoculated at 1 - 3 million cells/ ml, grown at 25°C and split every 2-4 days.

### RT-qPCR

3 - 5 million cells were lysed in 1 ml Trizol (ThermoFisher Scientific) and total RNA was purified using the NucleoSpin RNA kit (Macherey-Nagel). Reverse-transcription (RT) was performed using SuperScript IV and Random hexamers following the manufacturer’s instructions (ThermoFisher Scientific). A similar protocol was followed for whole larvae (20 L3 larvae) and larval imaginal discs (50 wing discs or 150 genital discs). Between 88 ng and 1 μg total RNA was used for the RT reactions. cDNA was amplified using Advanced Universal SYBRGreen Supermix (BioRad) and 0.5 μM of corresponding primers. With the exception of the *Abd-B* C promoter, primers were designed over intron boundaries to prevent amplification from genomic DNA using the online version of Primer3 software (https://primer3.ut.ee/). Primers were annealed at 55°C or 60°C and Real-time PCR was performed using a LightCycler 480 instrument (Roche). mRNA abundance was normalized to the *Gapdh2* and *Act42A* housekeeping genes using the ΔΔCt method (Livak and Schmittgen, 2001). Experiments were performed on biological duplicates (n=2). Primer sequences for RT-qPCR amplification are listed in Table S1.

### Calibrated RT-qPCR for absolute quantification of *Abd-B* isoforms

To determine absolute mRNA abundance of *Abd-B* isoforms, external calibration was performed using a plasmid containing all *Abd-B* isoform-specific RT-qPCR amplicons. The plasmid was assembled using the NEBuilder HiFi DNA Assembly kit (NEB) into a linearized pBlueScript backbone using larval cDNA as a template. Primer sequences for Gibson assembly are listed in Table S1. Primers were annealed at 55°C or 60°C and qPCR was performed using the SsoAdvanced Universal SYBR Green Supermix (BioRad) on a LightCycler 480 instrument (Roche). mRNA abundance for the different isoforms was normalized over the reference plasmid and a primer set to the common exon of all isoforms. Experiments were performed on biological duplicates (n=2). Primer sequences for RT-qPCR amplification are listed in Table S1.

### ChIP-seq and ChIP-qPCR

50 million S2, Sg4 or S3 cells were cross-linked in 1% formaldehyde for 10 min at room temperature. After quenching the cross-link reaction by adding 125 mM glycine, cells were lysed sequentially in buffers containing 10% Glycerol, 50 mM Tris-HCl pH 8.0, 140 mM NaCl, 1 mM EDTA, 0.5 % NP-40, 0.25 % Triton X-100 and 0.33X Complete Protease Inhibitors (Roche), followed by a buffer containing 10 mM Tris-HCl pH 7.5, 200 mM NaCl, 1 mM EDTA and 0.33X Complete Protease Inhibitors and a buffer containing 10 mM Tris-HCl pH 7.5, 100 mM NaCl, 1 mM EDTA, 0.25% SDS, 0.1% NaDeoxycholate and 1X Complete Protease Inhibitors. Chromatin was sheared using a Covaris S220 device with the following parameters: Peak Power: 110W, Duty Factor: 15%, Cycler/burst: 200, Duration: 1200 sec. After clarification by centrifugation, aliquots of sheared chromatin were kept at - 80°C.

For ChIP experiments, 3 μg of sonicated chromatin was diluted in a buffer containing 16 mM Tris-HCl pH 8.0, 167 mM NaCl, 1.2 mM EDTA, 0.01% SDS, 1.1% Triton X-100 and 1x Complete Protease Inhibitors (Roche) and precleared with Salmon Sperm DNA/Protein A Agarose beads (Merck-Millipore) for 30 min at 4°C. 10% of the sample was taken as input sample and the remainder was incubated with the specific antibody overnight at 4°C. The following antibodies were used: 2 μL H3K4me3 (07-473, Merck-Millipore) or 5 μg H3K27me3 (17-622, Merck-Millipore). Antibody-bound chromatin was isolated using Salmon Sperm DNA/Protein A Agarose beads, followed by sequential washing with Low Salt Immune Complex Wash Buffer (20 mM Tris-HCl pH 8.0, 150 mM NaCl, 2 mM EDTA, 0.1 % SDS and 1 % Triton X-100), High Salt Immune Complex Wash Buffer (20 mM Tris-HCl pH 7.5, 500 mM NaCl, 2mM EDTA, 0.1 % SDS, 1 % Triton X-100), LiCl Immune Complex Wash Buffer (10 mM Tris-HCl pH 7.5, 1 mM EDTA, 0.26 M LiCl, 2% NP-40 and Na Deoxycholate) and twice in 10mM Tris-HCl, 1mM EDTA. Chromatin was eluted using a 1% SDS, 0.1 mM NaHCO_3_ solution, followed by decrosslinking.

For ChIP-qPCR, experiments were performed using the SsoAdvanced Universal SYBR Green Supermix (BioRad) on a LightCycler 480 instrument (Roche). Enrichment was determined relative to Input using the ΔCt method. Experiments were performed on biological duplicates (n=2). Primer sequences for ChIP-qPCR amplification are listed in Table S1.

For ChIP-seq, Illumina sequencing libraries were prepared using the NEBNext Ultra II FS DNA library kit (NEB) and sequenced on the Illumina NextSeq 500 or 550 system (75 bp single end reads). ChIP-seq sequencing reads were filtered using FastQC filtering (https://www.bioinformatics.babraham.ac.uk/projects/fastqc/), followed by mapping onto the *Drosophila* genome (dm6) using bowtie2 (version 2.3.0) with filtering for multiple alignments (Langmead and Salzberg, 2012). Low quality reads (<30) and PCR duplicates were removed using SAMtools (Danecek *et al*., 2021). Bedgraphs were generated using BAMCoverage (Ramirez *et al*., 2016).

### CUT&Tag

CUT&Tag for H3K4me3 on WT and Fab-7 mutant genital discs was performed as described (Ahmad and Henikoff, 2020; Ahmad and Henikoff, 2021), with the following modifications. Larvae were harvested and washed in 1X PBS and then transferred to a buffer containing 20 mM HEPES pH 7.5, 150 mM NaCl, 0.5 mM Spermidine and 1X Complete Protease Inhibitors (Roche). 80 genital discs per genotype were dissected and collected in 200 μL of the same buffer, followed by addition of 10 μL Concanavalin A-coated beads suspension. After a 10 min incubation, the solution was removed and discs were incubated O/N at 4°C in 50 μL of 1% diluted H3K4me3 antibody (07-473, Merck-Millipore). After removal of the diluted H3K4me3 solution, discs were incubated for 40 minutes in 50 μL of 1% secondary antibody solution. For the CUT&Tag reaction, the discs were first incubated for 1 hour in 50 μL of home-made pAG-Tn5 (IJM, France). Discs were then washed with 50 μL of a buffer containing 20 mM HEPES pH 7.5, 300 mM NaCl, 0.5 mM Spermidine and 1X Complete Protease Inhibitors followed by tagmentation for 1 hour at 37°C in 50 μL of a buffer containing 10mM MgCl_2_, 20 mM HEPES pH 7.5, 300 mM NaCl, 0.5 mM Spermidine and 1X Complete Protease Inhibitors. The reaction was stopped by adding 50 μL of a buffer containing 0.17% SDS, 0.3 mg Proteinase K, 20 mM HEPES pH 7.5, 300 mM NaCl, 0.5 mM Spermidine and 1X Complete Protease Inhibitors and incubation for 1 hour at 58°C. DNA was purified using phenol-chloroform-isoamyl alcohol extraction and ethanol precipitation. Libraries were amplified using the NEBNext High-Fidelity 2X PCR Master Mix (NEB). CUT&Tag libraries were sequenced on the Illumina Nextseq 550 system (75 bp paired end reads) and analyzed as described for the ChIP-seq data.

### 4C-seq

Suitable 4C viewpoints were selected within or near the promoter of selected genes based on previously published criteria and genomic sequences obtained from the *Drosophila* dm6 release (Matelot and Noordermeer, 2016). PAGE-purified sequencing primers containing Illumina Tru-seq adapters and indexes (Eurogentec) are listed in Table S1.

4C-seq in cell lines was performed as published (Matelot and Noordermeer, 2016) with modifications: 50 million cells were cross-linked in 2% formaldehyde for 10 min at room temperature and lysed in a buffer containing 50 mM Tris-HCl pH 7.5, 150 mM NaCl, 5 mM EDTA, 0.5% NP-40, 1% Triton and 1X Complete Protease Inhibitors (Roche). After lysis, cell aliquots were stored at - 80°C. 4C-seq libraries were generated using DpnII (NEB) as first restriction enzyme and NlaIII (NEB) as the second restriction enzyme. Ligation reactions were performed using high concentration T4 DNA Ligase (Promega). For each viewpoint, 12 PCR reactions each containing 12 ng were performed using the Expand Long Template PCR System (Roche) with 30 cycles of amplification on a Thermocycler (Bio-Rad C1000). PCR reactions were pooled and purified using a PCR Clean up kit (Qiagen). Up to 23 viewpoints were mixed in equimolar ratio and sequenced on the Illumina NextSeq500 or 550 system (75 bp single end reads).

For 4C-seq in larval imaginal discs, a minimum of 1600 genital discs or 400 wing discs were dissected in cold 1X PBS over multiple sessions that lasted a maximum of 1 hour. Chromatin crosslinking and cell lysis were performed as for cell lines after dissection, with the addition of a second cell lysis step using a glass douncer with pestle. After lysis, cell aliquots were stored at - 80°C. After pooling of imaginal discs, the experimental procedure as outlined for cell lines was followed.

4C-seq sequencing reads were sorted, aligned, and translated to restriction fragments using the C4CTUS tool, which is a stand-alone version of the 4C-seq module within the former HTSstation online data analysis service (David *et al*., 2014) (C4CTUS is available at https://github.com/NoordermeerLab/c4ctus). Reads from different biological samples were demultiplexed using their Illumina Index and reads from different viewpoints were demultiplexed using the first 18 bases sequenced (viewpoint-specific primer). After removing the sequence of the viewpoint-specific primer, the remainder of the reads were mapped onto the *Drosophila* genome (dm6). Reads mapping to the viewpoint, the directly neighboring “undigested” fragment and fragments 2 kb up- and downstream were excluded during the procedure. Fragment counts were normalized per one million reads in a region within chr3R:4,661,427-8,999,228 (for viewpoints in the ANT-C) or chr3R:14,656,623-18,972,236 (for viewpoints in the BX-C). For visualizations, all tracks were smoothed using a running mean transformation over 11 consecutive valid restriction fragments.

### Hi-C

For Sg4 cells, Hi-C was performed on 15 million cells using the Arima Hi-C kit (Arima Genomics), following the manufacturer’s instructions. Hi-C material was sequenced on the Illumina NovaSeq 6000 system (150 bp paired end reads. Hi-C data from S2 R+ cells were obtained from the GEO repository GSE99107 (Szabo *et al*., 2018).

Hi-C reads were mapped to the *Drosophila* genome (dm6) using HiC-Pro (version 2.11.1) and bowtie2 (version 1.1.2) (Langmead and Salzberg, 2012; Servant *et al*., 2015). Default settings were used to remove duplicates, assign reads to their restriction fragments and filter for valid interactions. Hi-C matrices at 5kb resolution were generated from the valid interactions and normalized with the Iterative Correction and Eigenvector decomposition method (ICE) in the HiC-Pro tool. TADtool was used to call TAD boundaries with a window size of 51kb and a cutoff of 47 (S2 cells) or 75 (Sg4 cells) (Kruse *et al*., 2016).

### Single-cell RNA-seq

For single-cell RNA-seq experiments on cell lines, one million S2 and Sg4 cells were diluted in PBS (without MgCl_2_) and 0.04% BSA to a concentration of 1000 cells / μl. Reverse transcription and sequencing library preparation were done using the Chromium Next GEM Single Cell 3’ kit (10X Genomics), according to the manufacturer’s instructions. 3’-single-cell RNA-seq libraries were sequenced on the Illumina NextSeq 550 system (75 bp paired end reads).

For single-cell RNA-seq experiments on WT and Fab-7^1^ larval imaginal discs, pools of discs were dissected within 20 min in PBS (without MgCl_2_) with 0.04% BSA. To obtain roughly equivalent numbers of cells for each type of imaginal disc, the following pools of discs were combined: 4 eye/antenna discs, 8 leg 1 discs, 4 leg 2 discs, 4 leg 3 discs, 8 halter discs, 2 wing discs and 16 genital discs. Pools of discs were dissociated in 0.05% Trypsin EDTA (Thermofisher Scientific) for 8 minutes at 37°C and resuspended in PBS (without MgCl_2_) with 0.04% BSA to a concentration of approximately 1000 cells / μl. Reverse transcription and sequencing library preparation were done using the Chromium Next GEM Single Cell 5’ kit (10X Genomics), according to the manufacturer’s instructions. 5’ - singlecell RNA-seq libraries were sequenced on the Illumina NextSeq 550 system (75 bp paired end reads).

FastQ files were analyzed using the Cell Ranger software (10X Genomics, version 3.1.0), including alignment, filtering and quantitation of reads on the *Drosophila* genome (dm6) and generation of feature-barcode matrices. For the Sg4 and S2 cell lines, the clusters were obtained using k-means clustering (k = 3) using the Cell Ranger software. Further visualizations were generated using the Loupe browser (10X Genomics, version 4.2.0). For the imaginal discs, all downstream analyses were performed using the Seurat tool (version 4.0.1) (Hao *et al*., 2021). Cells with UMI count lower than 350 and mitochondrial genes above 5.5% were excluded from the analysis. After log-normalization, the vst method was used to select the top 2,000 variable features. WT and Fab-7^1^ imaginal disc cell data were then integrated using the dataset with the highest number of cells and the first 10 dimensions as a reference. Clustering was performed with the functions FindNeighbors and FindClusters, using the first 10 dimensions from PCA, resulting in the identification of 14 clusters. Identification of the clusters was done using the FindConservedmarkers function in Seurat. Cluster identities were assigned based on *Hox* gene expression for imaginal discs whereas for neurons, muscle and dead cells, the identities were assigned using the Flybase databases. Muscle cells and genital discs were initially assigned to two clusters each, which we merged for further analysis. All visualizations were performed using Seurat as well.

## Notes

### Competing Interest Statement

The authors have declared no competing interest.

