## Supplemental figures for "The *Drosophila* Fab-7 boundary element modulates *Abd-B* gene activity in the genital disc by guiding an inversion of collinear chromatin organization and alternative promoter use"

### FIGURE S1

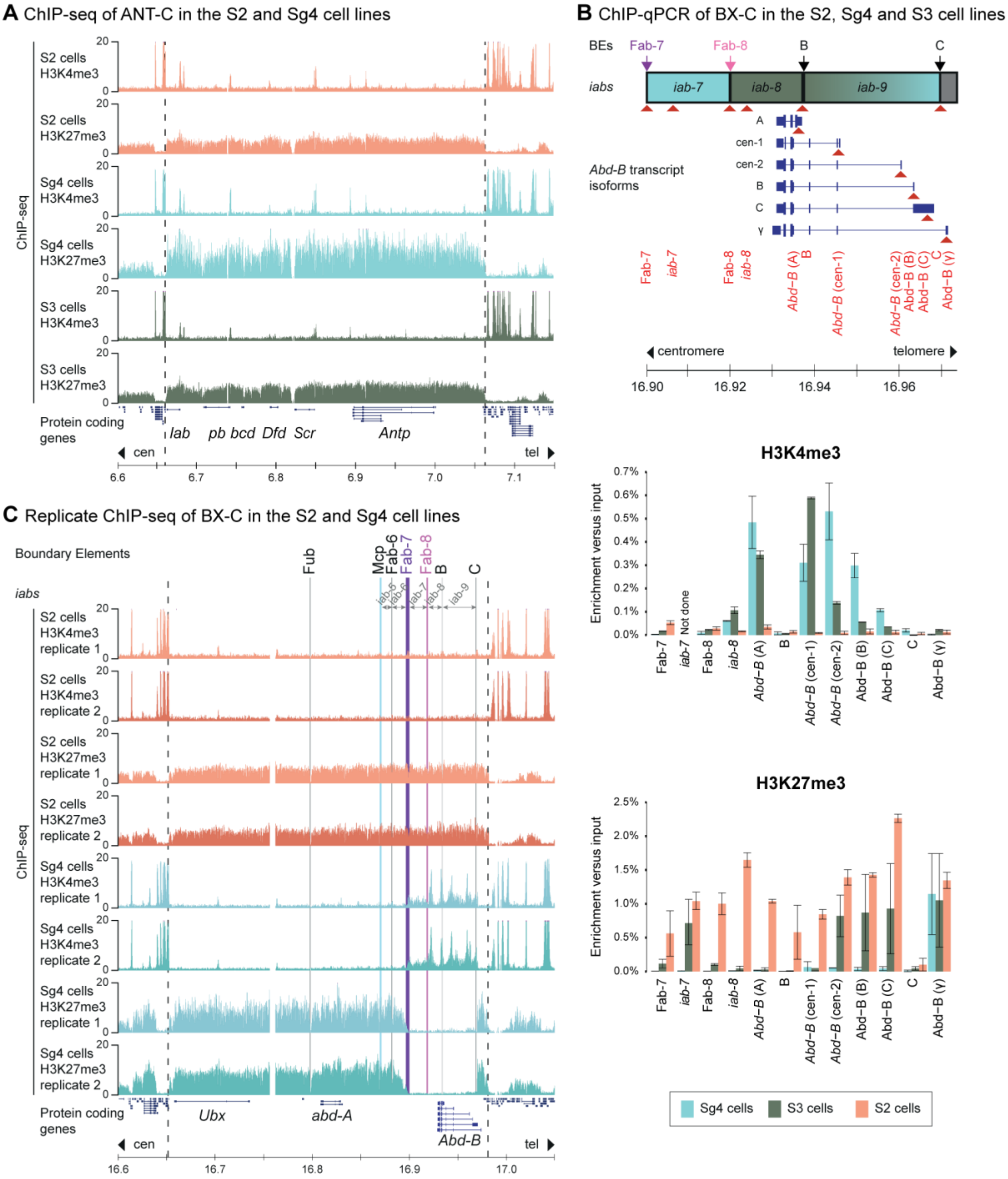
A ChIP-seq of ANT-C in the S2 and Sg4 cell lines
B ChIP-qPCR of BX-C in the S2, Sg4 and S3 cell lines
C Replicate ChIP-seq of BX-C in the S2 and Sg4 cell lines
ChIP-seq
S2 cells H3K4me3
S2 cells H3K27me3
Sg4 cells H3K4me3
Sg4 cells H3K27me3
S3 cells H3K4me3
S3 cells H3K27me3
Protein coding genes
lab pb bcd Dfd Scr Antp
cen
tel
BEs
Fab-7
Fab-8
iabs
iab-7
iab-8
iab-9
Abd-B transcript isoforms
centromere
telomere
H3K4me3
H3K27me3
Enrichment versus input
Not done
Sg4 cells
S3 cells
S2 cells
Boundary Elements
Fub
Mcp
Fab-6
Ubx
abd-A
Abd-B

**Figure S1: Histone modification landscape of the ANT-C and BX-C. Related to Figure 1.**

**A.** ChIP-seq data for the H3K4me3 and H3K27me3 histone marks in the S2, Sg4 and S3 cell lines at the ANT-C. The ANT-C is demarcated by dashed lines. Location of protein coding transcripts is indicated below.

**B.** ChIP-qPCR for H3K27me and H3K4me3 in the S2, S3 and Sg4 cell lines. Data are normalized over input. The position of analyzed regions within the *Abd-B cis*-regulatory domain is indicated above (red arrow heads). Data based on biological replicates (n = 2). Error bars indicate Standard Deviation.

**C.** ChIP-seq data for H3K4me3 and H3K27me3 at the BX-C in biological replicates for the S2 and Sg4 cell lines.

### FIGURE S2

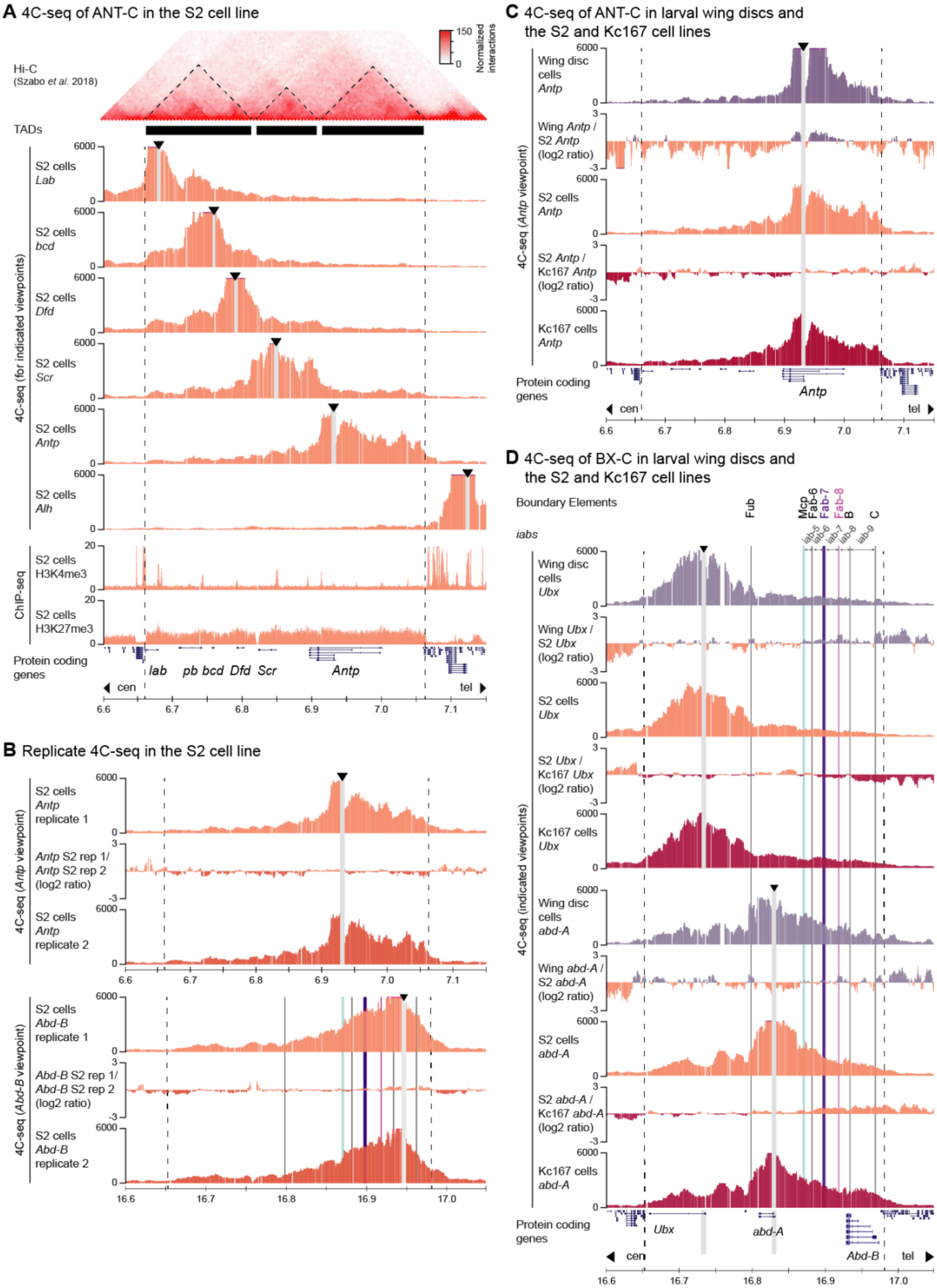

A 4C-seq of ANT-C in the S2 cell line
Hi-C
(Szabo et al. 2018)
150
0
Normalized interactions
TADs
4C-seq (for indicated viewpoints)
S2 cells Lab
S2 cells bcd
S2 cells Dfd
S2 cells Scr
S2 cells Antp
S2 cells Alh
ChIP-seq
S2 cells H3K4me3
S2 cells H3K27me3
Protein coding genes
lab pb bcd Dfd Scr Antp
cen
tel
B Replicate 4C-seq in the S2 cell line
4C-seq (Antp viewpoint)
S2 cells Antp replicate 1
Antp S2 rep 1/ Antp S2 rep 2 (log2 ratio)
S2 cells Antp replicate 2
4C-seq (Abd-B viewpoint)
S2 cells Abd-B replicate 1
Abd-B S2 rep 1/ Abd-B S2 rep 2 (log2 ratio)
S2 cells Abd-B replicate 2
C 4C-seq of ANT-C in larval wing discs and the S2 and Kc167 cell lines
4C-seq (Antp viewpoint)
Wing disc cells Antp
Wing Antp / S2 Antp (log2 ratio)
S2 cells Antp
S2 Antp / Kc167 Antp (log2 ratio)
Kc167 cells Antp
Protein coding genes
Antp
D 4C-seq of BX-C in larval wing discs and the S2 and Kc167 cell lines
Boundary Elements
Fub
Mcp
Fab-6
Fab-7
Fab-8
B
C
iabs
iab-5
iab-6
iab-7
iab-8
iab-9
4C-seq (indicated viewpoints)
Wing disc cells Ubx
Wing Ubx / S2 Ubx (log2 ratio)
S2 cells Ubx
S2 Ubx / Kc167 Ubx (log2 ratio)
Kc167 cells Ubx
Wing disc cells abd-A
Wing abd-A / S2 abd-A (log2 ratio)
S2 cells abd-A
S2 abd-A / Kc167 abd-A (log2 ratio)
Kc167 cells abd-A
Protein coding genes
Ubx abd-A Abd-B

**Figure S2: 3D chromatin organization of the ANT-C and BX-C in repressed cell types. Related to Figure 2.**

- **A.** Hi-C (top) and 4C-seq (middle) and ChIP-seq (bottom) data for the ANT-C in the S2 cell line. Hi-C data was reanalyzed from (Szabo *et al.*, 2018), with identified sub-domains indicated as black bars. 4C-seq data for viewpoints in the promoters of the *Lab, bcd, Dfd, Scr, Antp* and *Alh* genes is indicated in-between. H3K4me3 and H3K27 ChIP-seq data is indicated below. The ANT-C is demarcated by dashed lines. Location of protein coding transcripts is indicated below. Arrowheads indicate the positions of viewpoints.
- **B.** 4C-seq data in biological replicates for the S2 cell line for the indicated viewpoints. In-between, the log2 ratio of interactions is shown.
- **C.** 4C-seq data in wing disc cells and the S2 and Kc167 cell lines for the *Antp* viewpoint. In-between, the log2 ratio of interactions is shown.
- **D.** 4C-seq data in wing disc cells and the Sg4 and S3 lines for the *Ubx* and *abd-A* viewpoints. In-between, the log2 ratio of interactions is shown. The BX-C is demarcated by dashed lines, with known BEs highlighted by vertical bars. BEs and *iabs* relevant for *Abd-B* regulation are indicated on top. Location of protein coding transcripts is indicated below.

### FIGURE S3

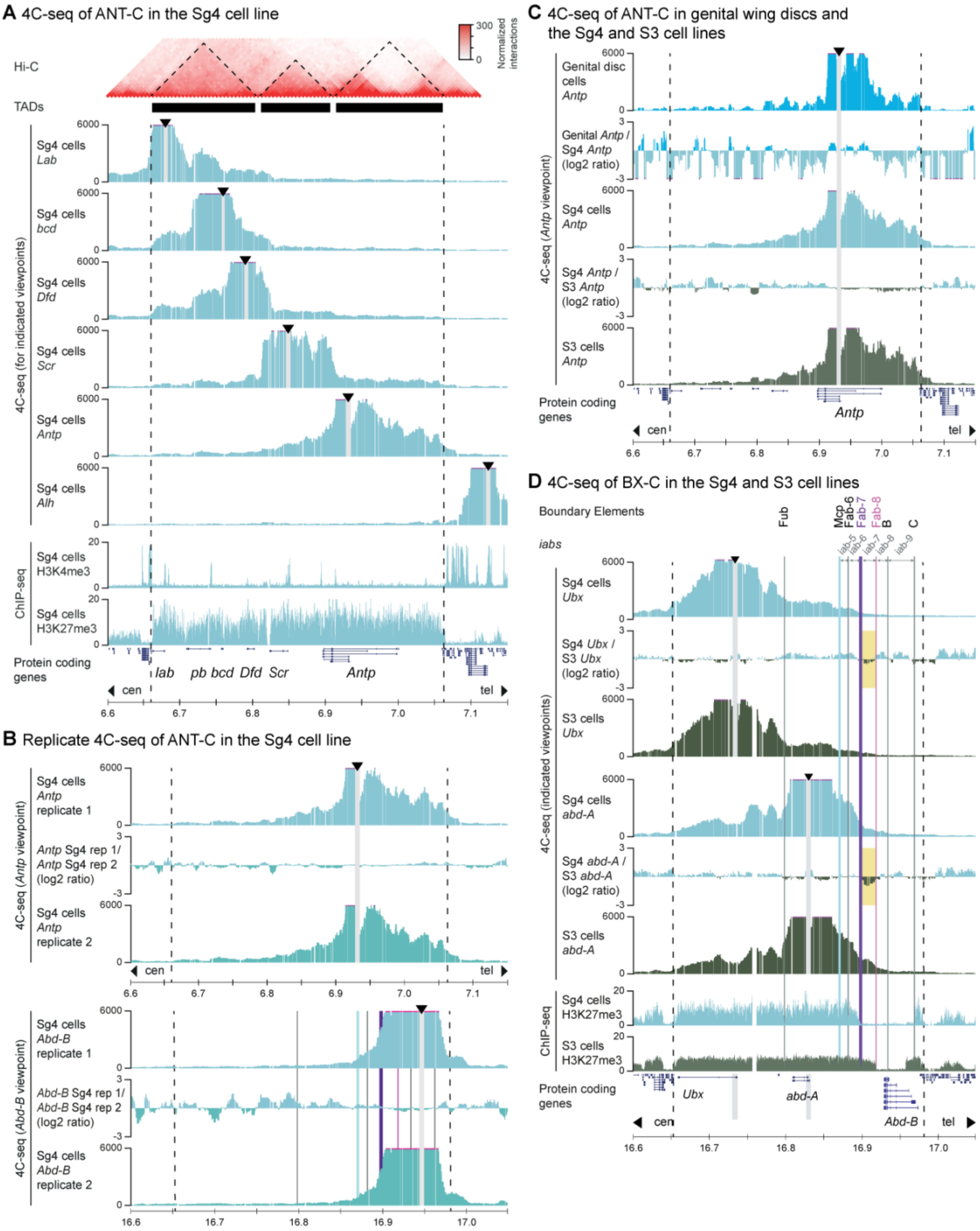


**Figure S3: 3D chromatin organization of the ANT-C and BX-C in cell types where *Abd-B* is active. Related to Figure 3.**

**A.** Hi-C (top) and 4C-seq (middle) and ChIP-seq (bottom) data for the ANT-C in the Sg4 cell line. Sub-domains as identified from Hi-C are indicated as black bars. 4C-seq data for viewpoints in the promoters of the *Lab*, *bcd*, *Dfd*, *Scr*, *Antp* and *Alh* genes is indicated in-between. H3K4me3 and H3K27 ChIP-seq data is indicated below. The ANT-C is demarcated by dashed lines. Location of protein coding transcripts is indicated below. Arrowheads indicate the positions of viewpoints.

**B.** 4C-seq data in biological replicates for the Sg4 cell line for the indicated viewpoints. In-between, the log2 ratio of interactions is shown.

**C.** 4C-seq data in genital disc cells and the Sg4 and S3 cell lines for the *Antp* viewpoint. In-between, the log2 ratio of interactions is shown.

**D.** 4C-seq data in the Sg4 and S3 cell lines for the *Ubx* and *abd-A* viewpoints. In-between, the log2 ratio of interactions is shown. The BX-C is demarcated by dashed lines, with known BEs highlighted by vertical bars. BEs and *iabs* relevant for *Abd-B* regulation are indicated on top. Location of protein coding transcripts is indicated below.

### FIGURE S4

**A** 4C-seq of BX-C in WT and Fab-7¹ wing disc cells

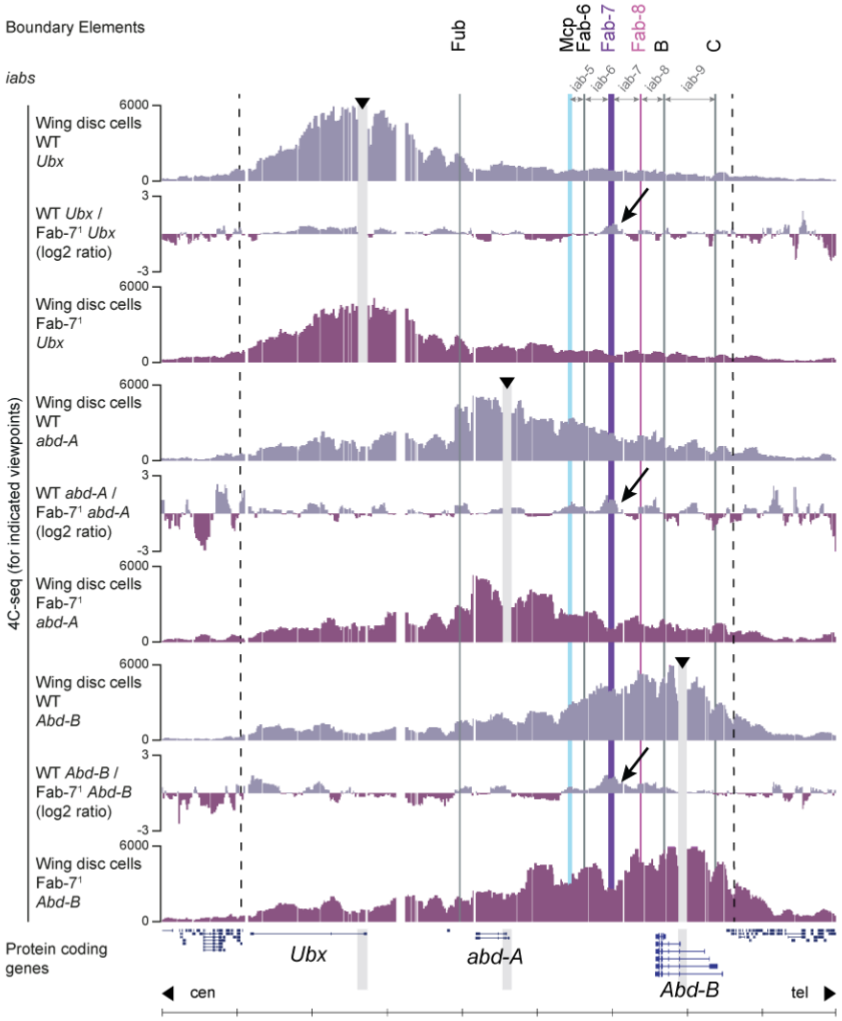


**B** 4C-seq of BX-C in WT and Fab-7¹ genital disc cells

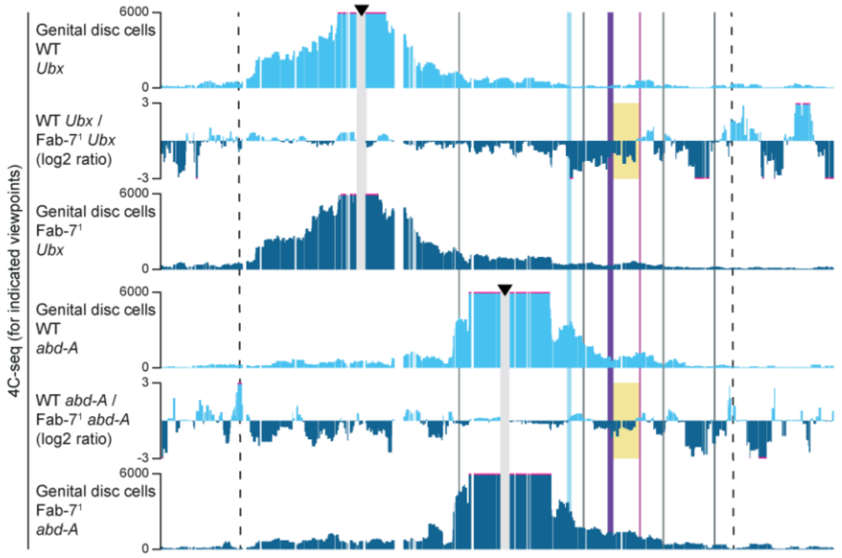

**Figure S4: 3D chromatin organization of the repressed BX-C in WT and Fab-7$^1$ deletion wing disc cells. Related to Figure 4.**

**A.** 4C-seq data in WT and Fab-7$^1$ deletion wing disc cells for the *Ubx*, *abd-A* and *Abd-B* viewpoints. In-between, the log2 ratio of interactions is shown. The BX-C is demarcated by dashed lines, with known BEs highlighted by vertical bars. BEs and *iabs* relevant for *Abd-B* regulation are indicated on top. Location of protein coding transcripts is indicated below. Arrowheads indicate the positions of viewpoints. Black arrows indicate the gain of signal in the WT cells where the Fab-7 element is present.

**B.** 4C-seq data in WT and Fab-7$^1$ deletion genital disc cells for the *Ubx* and *abd-A* viewpoints. In-between, the log(2) ratio of interactions is shown. The iab-7 where interactions are strongly increased in the Fab-7$^1$ deletion cells is highlighted with the yellow rectangle. Genomic coordinates and position of genes as in Figure S4A.

### FIGURE S5

**A** Number of UMIs per cell

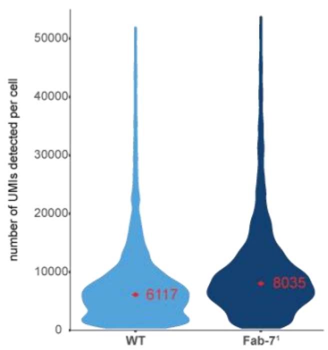


**B** Number of genes detected per cell

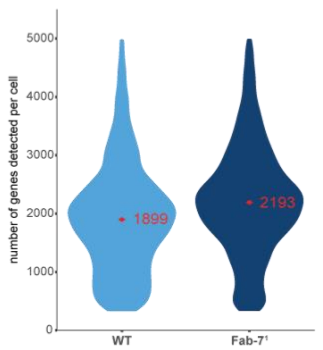


**C** UMAP projection of single-cell RNA-seq data for individual *Hox* genes

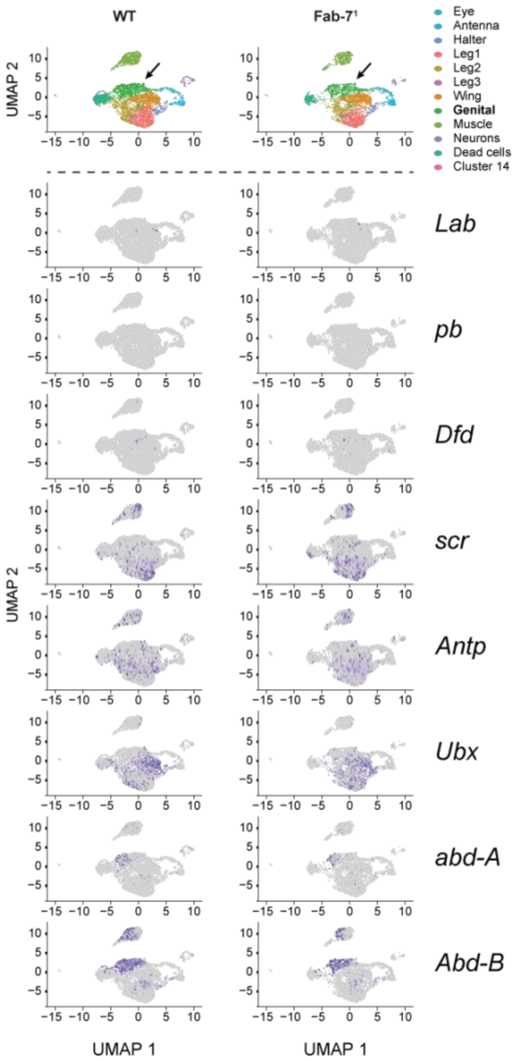


**D** Percentage of cells here *Hox* genes are (co-)detected

**Genital disc cells:**

| WT | *Lab* | *pb* | *Dfd* | *scr* | *Antp* | *Ubx* | *abd-A* | *Abd-B* |
|---|---|---|---|---|---|---|---|---|
| *Lab* | **1.2%** | 0.0% | 0.6% | 0.0% | 0.0% | 0.0% | 0.0% | 0.2% |
| *pb* | | **0.4%** | 0.1% | 0.0% | 0.0% | 0.1% | 0.2% | 0.3% |
| *Dfd* | | | **2.4%** | 0.1% | 0.0% | 0.2% | 0.2% | 0.6% |
| *scr* | | | | **9.0%** | 3.5% | 0.9% | 1.1% | 3.1% |
| *Antp* | | | | | **14.6%** | 3.6% | 1.5% | 5.3% |
| *Ubx* | | | | | | **16.0%** | 2.2% | 6.4% |
| *abd-A* | | | | | | | **21.2%** | 20.2% |
| *Abd-B* | | | | | | | | **65.2%** |

| Fab-7¹ | *Lab* | *pb* | *Dfd* | *scr* | *Antp* | *Ubx* | *abd-A* | *Abd-B* |
|---|---|---|---|---|---|---|---|---|
| *Lab* | **2.4%** | | | | | | | |
| *pb* | 0.1% | **0.6%** | | | | | | |
| *Dfd* | 0.3% | 0.1% | **3.1%** | | | | | |
| *scr* | 0.0% | 0.1% | 0.9% | **14.9%** | | | | |
| *Antp* | 0.1% | 0.0% | 0.1% | 6.9% | **18.5%** | | | |
| *Ubx* | 0.3% | 0.1% | 0.3% | 1.8% | 5.0% | **18.9%** | | |
| *abd-A* | 0.0% | 0.1% | 0.3% | 1.2% | 2.0% | 2.2% | **17.7%** | |
| *Abd-B* | 0.1% | 0.3% | 0.5% | 4.3% | 6.5% | 6.6% | 17.5% | **49.0%** |

**Wing disc cells:**

| WT | *Lab* | *pb* | *Dfd* | *scr* | *Antp* | *Ubx* | *abd-A* | *Abd-B* |
|---|---|---|---|---|---|---|---|---|
| *Lab* | **0.2%** | 0.0% | 0.0% | 0.0% | 0.0% | 0.0% | 0.0% | 0.0% |
| *pb* | | **0.0%** | 0.0% | 0.0% | 0.0% | 0.0% | 0.0% | 0.0% |
| *Dfd* | | | **0.6%** | 0.0% | 0.0% | 0.2% | 0.0% | 0.1% |
| *scr* | | | | **5.4%** | 1.7% | 2.9% | 0.0% | 0.2% |
| *Antp* | | | | | **20.2%** | 8.4% | 0.2% | 0.8% |
| *Ubx* | | | | | | **59.8%** | 0.5% | 2.8% |
| *abd-A* | | | | | | | **0.8%** | 0.5% |
| *Abd-B* | | | | | | | | **4.7%** |

| Fab-7¹ | *Lab* | *pb* | *Dfd* | *scr* | *Antp* | *Ubx* | *abd-A* | *Abd-B* |
|---|---|---|---|---|---|---|---|---|
| *Lab* | **0.7%** | | | | | | | |
| *pb* | 0.0% | **0.0%** | | | | | | |
| *Dfd* | 0.4% | 0.0% | **1.0%** | | | | | |
| *scr* | 0.4% | 0.0% | 0.5% | **7.1%** | | | | |
| *Antp* | 0.3% | 0.0% | 0.1% | 2.9% | **25.1%** | | | |
| *Ubx* | 0.5% | 0.0% | 0.3% | 3.1% | 6.4% | **44.8%** | | |
| *abd-A* | 0.0% | 0.0% | 0.0% | 0.0% | 0.5% | 1.0% | **1.2%** | |
| *Abd-B* | 0.3% | 0.0% | 0.1% | 1.0% | 2.7% | 5.2% | 0.9% | **8.7%** |

**E** t-SNE projection of single-cell RNA-seq data for *Abd-B* and *abd-A* in S2 and Sg4 cells

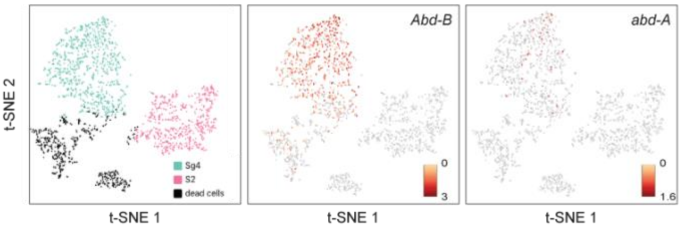


**F** Detection of *Abd-B* isoforms in genital disc cells

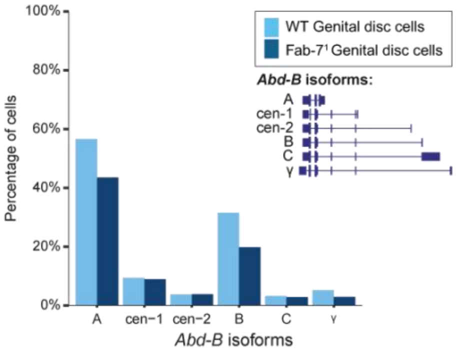

**Figure S5: BX-C gene and promoter activity in individual cells. Related to Figure 5.**

- **A.** Violin plots showing the number of detected Unique Molecular Identifiers (UMIs) per cell in single-cell RNA-seq data from pools of WT and Fab-$7^1$ deletion imaginal disc cells. Red diamonds and numbers indicate the median.
- **B.** Violin plots showing the number of detected genes per cell in single-cell RNA-seq data from pools of WT and Fab-$7^1$ deletion imaginal disc cells. Red diamonds and numbers indicate the median.
- **C.** UMAP projections of single-cell RNA-seq data from pools of WT and Fab-$7^1$ deletion imaginal disc cells. On the top, the identified clusters are indicated. On the bottom, the presence of mRNA from individual *Hox* genes (purple) within the pool of discs is indicated.
- **D.** Tables showing the percentage of cells where mRNA from individual and pairs of *Hox* genes is detected in genital disc and wing disc cells.
- **E.** t-SNE projections of 3'- single-cell RNA-seq data from the S2 and Sg4 cell lines. On the left, the identified clusters are indicated. On the right, the presence of *Abd-B* or *abd-A* mRNA is indicated.
- **F.** Histogram showing the percentage of cells where mRNA for each *Abd-B* isoform is detected in genital disc cells (percentages within the entire population of genital disc cells, including those where no *Abd-B* mRNA is detected).
