## Supplemental Table S1 for "The *Drosophila* Fab-7 boundary element modulates *Abd-B* gene activity in the genital disc by guiding an inversion of collinear chromatin organization and alternative promoter use"

### Table S1: Primer sequences

#### RT-qPCR

| *Target* | *Sequence* |
|---|---|
| *Abd-B* Forward<br>*Abd-B* Reverse | GCACATCGACGTCCAGCTAT<br>GCTCCCACGGATAATCCACC |
| *abd-A* Forward<br>*Abd-A* Reverse | GCTTACAGACTGGATGGGAA<br>GGTTGAAGTGAAACTCCTTCTC |
| *Ubx* Forward<br>*Ubx* Reverse | TCGCAGGTAAGAGATACTCAG<br>GATTCGTGTGGAACTCCTTC |
| *Antp* Forward<br>*Antp* Reverse | AGATTAGGCTACGCAACTGTAC<br>GTGATCGTGACTGTGACTCTG |
| *Scr* Forward<br>*Scr* Reverse | CGTGGATGAAGCGAGTAC<br>GTAGCGGTTGAAGTGGAAC |
| *Dfd* Forward<br>*Dfd* Reverse | GCGAAGAAACCCACCAAG<br>CCGATACTCTCCAAACTGTC |
| *Act42A* Forward<br>*Act42A* Reverse | AAGTGTGTGCAGCGGATAACT<br>GTTGTCGACCACTAAAGCTGC |
| *Gapdh2* Forward<br>*Gapdh2* Reverse | CCCGAGTTTTCGCCCATAGA<br>CCGATGCGACCAAATCCATTG |

#### Calibrated RT-qPCR

| *Promoter* | *Target* | *Sequence* |
|---|---|---|
| Common exon | *Abd-B* 1 Forward<br>*Abd-B* 1 Reverse | GCACATCGACGTCCAGCTAT<br>GCTCCCACGGATAATCCACC |
| Promoter A<br>(α or m) | *Abd-B* A Forward<br>*Abd-B* A Reverse | GTGTTTCTCCCTCCAGTGTTAGT<br>GAGGTGTCCGTCTCGAATCC |
| Promoter cen-1 | *Abd-B* cen-1 Forward<br>*Abd-B* cen-1 Reverse | TAATGACGGCTACCCAACGG<br>CAGTTATTTGGGTAGTGCTTTTGGT |
| Promoter cen-2 | *Abd-B* cen-2 Forward<br>*Abd-B* cen-2 Reverse | CCAGAAACCGCAGATTTGAGC<br>CCGGTTTGCTCACTTCCAGT |
| Promoter B | *Abd-B* B Forward<br>*Abd-B* B Reverse | ATCAGACTTGCAGGTCACGTAT<br>TGCTTTTGGTCAAAATCGCTGA |
| Promoter C | *Abd-B* C Forward<br>*Abd-B* C Reverse | AGGAGCTTATGCGAATGTCTGT<br>CAGCTACCAAAACTCAATGCCA |
| Promoter γ | *Abd-B* γ Forward<br>*Abd-B* γ Reverse | TTCGGAAGATTGTATTTGTGCGG<br>CCGGTTTGCTCACTTCCAGT |

**ChIP-qPCR**

| ***Target*** | ***Sequence*** |
|---|---|
| Fab-7 Forward<br>Fab-7 Reverse | GCTCACTAACACATAGATAA<br>CTCTCTTGCTTACCAATAC |
| *iab-7* Forward<br>*iab-7* Reverse | GTTGACTAGAACCCAAATC<br>CCACACTCATCGTTATTC |
| Fab-8 Forward<br>Fab-8 Reverse | GGTTGGTCTGCATATTG<br>CGGATTTCTGCTTTCTG |
| *iab-8* Forward<br>*iab-8* Reverse | CGGCGATAATGTCTTTC<br>GGGCACATCTTTCTTATC |
| *Abd-B* (A) Forward<br>*Abd-B* (A) Reverse | GGCTGTTGTTGTAGTTG<br>CGTCAGTGTAGATTTGTG |
| B Forward<br>B Reverse | TGTGACTTGGAAGAGAG<br>GTGTACGTGTGCATAAC |
| *Abd-B* (cen-1) Forward<br>*Abd-B* (cen-1) Reverse | GTGATAAGACCAAAGGAATA<br>CGGGCCGATATTTAATG |
| *Abd-B* (cen-2 Forward<br>*Abd-B* (cen-2) Reverse | GGCTGCTTCATGTTTAC<br>GCGGCCAAGTTATTAAG |
| *Abd-B* (B) Forward<br>*Abd-B* (B) Reverse | GCGGTACTAGTGCTTTA<br>CCGAGTATTTACCACTTTAC |
| *Abd-B* (C) Forward<br>*Abd-B* (C) Reverse | CAATGCCAAACAAAGTATC<br>GAGCTTATGCGAATGTC |
| C Forward<br>C Reverse | TGCGCAGAGTCTTATAC<br>CTAGTGAACAGACCATCTA |
| *Abd-B* (γ) Forward<br>*Abd-B* (γ) Reverse | AACAACCGCACAAATAC<br>AACCGAGAACCCATTAG |
| Mcp Forward<br>Mcp Reverse | CTGTTGGAAACGCATTA<br>TGTGTAAGGAGGAAGAC |
| Fab-6 Forward<br>Fab-6 Reverse | TTGCCAGATAAATCCAAGAT<br>GCCGATTAAAGCAGTCC |

**4C-seq**

Color coding: Illumina adaptor sequence - index - target specific sequence.

| ***Target*** | ***Sequences*** |
|---|---|
| *Abd-B* promoter cen-1 | **Forward**:<br>AATGATACGGCGACCACCGAGATCTACACTCTTTCCCTACACGA CGCTCTTCCGATCTggtaatatggcaatcagcctc<br>**Reverse – index 1:**<br>CAAGCAGAAGACGGCATACGAGATCGTGATGTGACTGGAGTTCA GACGTGTGCTCTTCCGATCTaaagtacttactgaaccactctc<br>**Reverse – index 2:**<br>CAAGCAGAAGACGGCATACGAGATACATCGGTGACTGGAGTTCA GACGTGTGCTCTTCCGATCTaaagtacttactgaaccactctc |
| *iab-7* | **Forward:**<br>AATGATACGGCGACCACCGAGATCTACACTCTTTCCCTACACGA CGCTCTTCCGATCTcacctgctatcccaaagttc<br>**Reverse – index 1:**<br>CAAGCAGAAGACGGCATACGAGATCGTGATGTGACTGGAGTTCA GACGTGTGCTCTTCCGATCTggccgacttttgaatttgttt<br>**Reverse – index 2:**<br>CAAGCAGAAGACGGCATACGAGATACATCGGTGACTGGAGTTCA GACGTGTGCTCTTCCGATCTggccgacttttgaatttgttt |
| *abd-A* promoter | **Forward:**<br>AATGATACGGCGACCACCGAGATCTACACTCTTTCCCTACACGA CGCTCTTCCGATCTcggcttattgtcacttaaatgtta<br>**Reverse – index 1:**<br>CAAGCAGAAGACGGCATACGAGATCGTGATGTGACTGGAGTTCA GACGTGTGCTCTTCCGATCTatggcgcaatacaaaagcc<br>**Reverse – index 2:**<br>CAAGCAGAAGACGGCATACGAGATACATCGGTGACTGGAGTTCA GACGTGTGCTCTTCCGATCTatggcgcaatacaaaagcc |
| *Ubx* promoter | **Forward:**<br>AATGATACGGCGACCACCGAGATCTACACTCTTTCCCTACACGA CGCTCTTCCGATCTatccgccaaaaatcgcaga<br>**Reverse – index 1:**<br>CAAGCAGAAGACGGCATACGAGATCGTGATGTGACTGGAGTTCA GACGTGTGCTCTTCCGATCTcgccgcgggaaattcatc<br>**Reverse – index 2:**<br>CAAGCAGAAGACGGCATACGAGATACATCGGTGACTGGAGTTCA GACGTGTGCTCTTCCGATCTcgccgcgggaaattcatc |
| *Alh* promoter | **Forward:**<br>AATGATACGGCGACCACCGAGATCTACACTCTTTCCCTACACGA CGCTCTTCCGATCTaactagacccgcactctcc<br>**Reverse – index 1:**<br>CAAGCAGAAGACGGCATACGAGATCGTGATGTGACTGGAGTTCA GACGTGTGCTCTTCCGATCTtggaatacgggtgggtgt<br>**Reverse – index 2:**<br>CAAGCAGAAGACGGCATACGAGATACATCGGTGACTGGAGTTCA GACGTGTGCTCTTCCGATCTtggaatacgggtgggtgt |

| | |
|---|---|
| *Antp* promoter | **Forward:**<br>AATGATACGGCGACCACCGAGATCTACACTCTTTCCCTACACGA CGCTCTTCCGATCTgagatggggatgtggttg<br>**Reverse – index 1:**<br>CAAGCAGAAGACGGCATACGAGATCGTGATGTGACTGGAGTTCA GACGTGTGCTCTTCCGATCTattgtggtgtaaaagtcggt<br>**Reverse – index 2:**<br>CAAGCAGAAGACGGCATACGAGATACATCGGTGACTGGAGTTCA GACGTGTGCTCTTCCGATCTattgtggtgtaaaagtcggt |
| *Scr* promoter | **Forward:**<br>AATGATACGGCGACCACCGAGATCTACACTCTTTCCCTACACGA CGCTCTTCCGATCTcacgcgcaatttgttaca<br>**Reverse – index 1:**<br>CAAGCAGAAGACGGCATACGAGATCGTGATGTGACTGGAGTTCA GACGTGTGCTCTTCCGATCTattttaagcgctttggaaca<br>**Reverse – index 2:**<br>CAAGCAGAAGACGGCATACGAGATACATCGGTGACTGGAGTTCA GACGTGTGCTCTTCCGATCTattttaagcgctttggaaca |
| *Dfd* promoter | **Forward:**<br>AATGATACGGCGACCACCGAGATCTACACTCTTTCCCTACACGA CGCTCTTCCGATCTtagtaccgtcccctctttac<br>**Reverse – index 1:**<br>CAAGCAGAAGACGGCATACGAGATCGTGATGTGACTGGAGTTCA GACGTGTGCTCTTCCGATCTgttccatatgtgagcggtat<br>**Reverse – index 2:**<br>CAAGCAGAAGACGGCATACGAGATACATCGGTGACTGGAGTTCA GACGTGTGCTCTTCCGATCTgttccatatgtgagcggtat |
| *Bcd* promoter | **Forward:**<br>AATGATACGGCGACCACCGAGATCTACACTCTTTCCCTACACGA CGCTCTTCCGATCTgacgatacctcacagccc<br>**Reverse – index 1:**<br>CAAGCAGAAGACGGCATACGAGATCGTGATGTGACTGGAGTTCA GACGTGTGCTCTTCCGATCTccgaatcttgcaatattactttca<br>**Reverse – index 2:**<br>CAAGCAGAAGACGGCATACGAGATACATCGGTGACTGGAGTTCA GACGTGTGCTCTTCCGATCTccgaatcttgcaatattactttca |
| *Lab* promoter | **Forward:**<br>AATGATACGGCGACCACCGAGATCTACACTCTTTCCCTACACGA CGCTCTTCCGATCTaattggatgtttcaaaacattttaag<br>**Reverse – index 1:**<br>CAAGCAGAAGACGGCATACGAGATCGTGATGTGACTGGAGTTCA GACGTGTGCTCTTCCGATCTaatagtagacaactcttcgcc<br>**Reverse – index 2:**<br>CAAGCAGAAGACGGCATACGAGATACATCGGTGACTGGAGTTCA GACGTGTGCTCTTCCGATCTaatagtagacaactcttcgcc |
